# Single-Trial Characterisation of Frontal Theta and Parietal Alpha Oscillatory Episodes during Spatial Navigation in Humans

**DOI:** 10.1101/2024.10.17.618670

**Authors:** Mireia Torralba Cuello, Angela Marti-Marca, Márta Szabina Pápai, Salvador Soto-Faraco

**Author notes:** **CORRESPONDING AUTHOR:** Mireia Torralba Cuello.

## Abstract

Theoretical proposals and empirical findings both highlight the relevance of theta brain oscillations in human Spatial Navigation. However, whilst the general assumption is that the relevant theta band activity is purely oscillatory, most empirical studies fail to disentangle oscillatory episodes from wide band activity. In addition, experimental approaches often rely on averaged activity across trials and subjects, disregarding moment-to-moment fluctuations in Theta activity, contingent on key aspects of the task. Here, we used novel oscillation detection approaches to investigate the dynamics of theta and alpha episodes in human subjects performing a spatial navigation task in a Virtual Reality environment, resolved at single-trial level. The results suggest that bouts of theta oscillatory activity are related to task difficulty and access to previously encoded information, across different timescales. alpha episodes, instead, seem to anticipate successful navigational decisions and could be related to shifts in internal attention.

## INTRODUCTION

Spatial navigation, an essential skill for many animal species including humans, is a complex ability that draws on multiple basic cognitive processes, such as memory, attention, proprioception, executive control, and decision making (Albert et al., 1999; Do et al., 2021; Eichenbaum, 2017; Parra-Barrero et al., 2023; Patai & Spiers, 2021). Spatial navigation capacity tends to decline with age, leading to a loss of autonomy and life quality in older adults (Burns, 1999; Lopez et al., 2018). Furthermore, topographical disorientation characterises some age-related conditions such as mild cognitive impairment (MCI) and Alzheimer disease (AD), and has been suggested as a potential biomarker of cognitive decline (Plácido et al., 2022). Due to its relevance, the cognitive and physiological bases of spatial navigation have been thoroughly investigated in non-human (Buzsáki, 2005; O’Keefe & Conway, 1978) and human animals (Epstein et al., 2017). However, whereas animal research and invasive methods in humans have highlighted the great relevance of episodes of oscillatory neural activity on a single-trial basis (Caplan et al., 2001; Jacobs et al., 2010; Nardin et al., 2023; Schmidt et al., 2009; Shin & Talnov, 2001), non-invasive human research has been more limited in this respect. Here, we build on recent advances in oscillation detection methods to characterise oscillatory episodes on human neural activity during learning and recalling paths in a virtual reality environment, using scalp electroencephalography (EEG).

Electrophysiological studies of spatial navigation in rodents have revealed the importance of hippocampal theta (3 to 8 Hz) oscillatory episodes (Buzsáki, 2005; Hasselmo & Stern, 2014; Jacobs, 2014; Winson, 1978), thought to help organise spatial information into cognitive maps that allow to integrate external information for flexible behaviour (O’Keefe & Nadel, 1978). More recent proposals have suggested that the role of the hippocampus is, more generally, to organise external cues into cognitive maps that can be accessed through memory. That is, cognitive maps include, but are not restricted to, spatial representations (Buzsáki, 2005; Eichenbaum, 2017; Epstein et al., 2017), an interpretation that harkens back to the concept of cognitive map originally introduced by Tolman (1948).

In humans, spatial navigation was traditionally assessed by “pen-and-paper” tests (Cogné et al., 2017), though the use of Virtual Reality (VR) is currently a popular approach (Sánchez-Escudero et al., 2024). VR environments can produce close-to-realistic immersive experiences whilst allowing for experimental control (Diersch & Wolbers, 2019). Furthermore, VR has been proposed as a valuable tool for assessing navigational disorders and as a potential device for interventions (Cogné et al., 2017), given that spatial navigation performance in VR environments and in real life are highly correlated (Cushman et al., 2008; Diersch & Wolbers, 2019). For example, age-related decline in spatial navigation skills in VR tasks seems to be comparable to real life (Cushman et al., 2008).

On the on the hand, non-invasive neuroimaging during spatial navigation in VR environments allows for the evaluation of neural correlates in humans during realistic navigation tasks, enabling the analysis of increasingly sophisticated patterns using oscillatory episode modelling (Caplan et al., 2001), source localization (Bauer et al., 2021), and multivariate analysis (Do et al., 2020; Zhu et al., 2022). Tasks like the Morris water maze (Thornberry et al., 2023), T-junction mazes (Bischof & Boulanger, 2003; Caplan et al., 2001), navigation through virtual cities (Araújo et al., 2002; Bischof & Boulanger, 2003), or real environments (Zhu et al., 2022), are some of the scenarios used in the study of human spatial navigation. Given that the brain operates through online computations, it is important to address relevant information contained in moment-to-moment variability as opposed to averages. At present, rich datasets and knowledge from neuroimaging studies make it possible to investigate, at single-trial level, neural dynamics in relation to the instantaneous navigational state. For instance single-trial analysis of phase precession showed that the phase span of precession is reduced when single-trial is analysed compared to pooled data (Schmidt et al., 2009).

One the other hand, invasive approaches using intracranial recordings of electrophysiological activity have allowed to link hippocampal theta to spatial navigation in humans, similar to earlier findings in other animal models (Bohbot et al., 2017; Ekstrom et al., 2005). In addition, fMRI and invasive electrophysiological studies suggest that an extended network of brain areas is involved in spatial navigation, including, but not limited to the hippocampus, parietal cortex and prefrontal cortex (Baumann & Mattingley, 2021; Caplan et al., 2001; Ekstrom et al., 2017; Hyman et al., 2005; Kahana et al., 1999; Patai & Spiers, 2021). Some of these studies show that it is possible to extract spatial navigation-related electrophysiological activity from scalp electroencephalography (EEG), which allows subjects to move freely, bringing the study of spatial navigation closer to near-realistic scenarios. These EEG studies have predominantly highlighted the relevance of oscillatory frequencies in the theta range (3-8 Hz) and, to a lesser extent, in alpha (8-14 Hz). It is important to clarify that invasively recorded hippocampal theta and theta in non-invasive recordings collected at fronto-medial sites most likely have different origins (Mitchell et al., 2008), although empirical findings in rodents (M. W. Jones & Wilson, 2005; Nardin et al., 2023; Siapas et al., 2005; Young & McNaughton, 2009) and humans (Gallinat et al., 2006) support a link between theta at fronto-medial locations extracted from non-invasive recordings and hippocampal theta.

The amplitude of theta oscillations (commonly observed in frontal areas in non-invasive electrophysiological recordings), has been reported to increase during movement compared to standing still (Liang et al., 2018a), during navigation compared to non-navigation periods (Araújo et al., 2002), in difficult compared to easy spatial navigation tasks (Caplan et al., 2001; Kahana et al., 1999), during free exploration compared to guided navigation (Chrastil et al., 2022), when subjects approach a target (Bauer et al., 2021), collide with (invisible) walls in virtual mazes (Gehrke & Gramann, 2021), must re-calculate paths (Do et al., 2021), and after a navigational decision (Bischof & Boulanger, 2003). Some studies have reported increased theta activity specifically after a successful navigational decision (Bauer et al., 2021; Chrastil et al., 2022; White et al., 2012), although not all studies have observed this correlation (Caplan et al., 2001). In summary, many studies report increases in theta oscillations during relevant spatial navigation events (although some other studies have also reported decreases in theta activity, associated with spatial learning (Thornberry et al., 2023)).

In addition to theta, non-invasive electrophysiological studies in humans have found correlates between spatial navigation and alpha oscillatory activity (8 to 14 Hz). Usually, alpha oscillatory activity is observed in parietal and occipital areas and, it has been reported to increase or decrease in different contexts. For example, alpha desynchronization has been suggested to reflect switches between allocentric and egocentric reference frames (Do et al., 2021). Processing of spatial navigation-relevant sensory information also results in alpha decreases during information gathering (Delaux et al., 2021), after collision with invisible obstacles (Gehrke & Gramann, 2021) or during navigation compared to non-navigation epochs (Araújo et al., 2002).

However, some authors associate posterior alpha desynchronization with visual flow, rather than specifically with spatial navigation operations (Liang et al., 2018a). Though less frequently reported, increases in alpha activity have also been observed, such as in free compared to guided navigation (Chrastil et al., 2022), or when walking away from invisible boundaries in a virtual maze (Gehrke & Gramann, 2021).

According to the studies briefly discussed above, there is evidence of the involvement of theta and alpha oscillations in various aspects of spatial navigation such as path difficulty, familiarity, encoding success, or navigation strategy. Yet, their specific role is far from clear. First, most of these past studies focus on mean activity across subjects and/or trials, therefore disregarding moment-to-moment online modulations of theta /alpha activity at different stages of the task (though see, Gehrke & Gramann, 2021; Zhu et al., 2022, for exceptions). This is particularly relevant given the importance of trial-level modulations in animal findings, and the effort that researchers put into measuring spatial navigation using realistic (and therefore complex) navigation paradigms. It is, therefore, crucial to evaluate the data with analytic approaches that address the neural correlates of navigation at a finer level of detail. Single-trial methods like linear mixed models are optimal for the study of neuroimaging data for several reasons, for instance, they are robust to unbalanced and incomplete data (Brown, V. A., 2021), allow for the modelling of individual variability, explicitly, without the need of aggregated information, their outcomes can be generalised beyond the sampled population (Boisgontier & Cheval, 2016). Second, despite many spatial navigation theories attributing significance to the oscillatory nature of theta activity, it is not always clear that the neural activity observed is really oscillatory. It is fundamental to disentangle transient episodes of oscillatory activity from background, aperiodic, neural activity (referred to sometimes as background 1/f activity), as they are usually conflated in the EEG signal (Gerster et al., 2022). Conventional time-frequency methods based on trial averages fail at this, as differences in the power of a certain frequency band can be due not only to differences in oscillatory activity but also to background activity (Donoghue et al., 2022), and estimates of peak (dominant) frequency can be strongly biased by aperiodic activity (Samaha & Cohen, 2022). Using averages of spectral estimates across time windows can be problematic in situations in which the stationarity of the data within the time window of interest is not met (S. R. Jones, 2016). Moreover, the accumulation of high power transient events across trials can produce the illusion of sustained oscillations in the aggregated data (S. R. Jones, 2016). Aggregated measures complicate the interpretability of the results, as increases in power at specific time-points can result both from increases in the “amplitude” of the signal and from an increase in the repeatability of oscillatory activity (Kosciessa et al., 2020). Moreover, the duration of oscillatory episodes can provide relevant information about the underlying neural processes generating the patterns of observed activity (van Ede et al., 2018).

Recent advances, based on the Better Oscillation detection method (BOSC) originally developed by Caplan et al. (2001) for intracranial EEG, now allow for the extraction of oscillatory episodes in scalp EEG data (Du et al., 2023; Liang et al., 2018). This approach allows the researcher to extract time-resolved, single-trial information of purely oscillatory activity, capturing transient episodes during spatial navigation in non-invasive data. So far, prior studies that have focused on oscillatory episodes either with invasive or non-invasive EEG (Caplan et al., 2001; Du et al., 2023; Kahana et al., 1999; Liang et al., 2018b), relied on simplified designs to satisfy a specific hypothesis-driven question (for instance, long vs. short mazes or correct vs. incorrect decisions). In the present study, we hypothesise that theta and alpha oscillatory activity encode relevant information related to spatial navigation and, therefore, we predict that oscillatory episodes at each of the frequency bands can be related to different aspects of the spatial navigation task. We will investigate modulations of the abundance of theta and alpha oscillatory episodes at single-trial level as humans perform a navigational memory task^1^ through a realistic virtual environment presented by means of a Head Mounted Display. Movement (translation and rotation) are simulated but controlled by the subjects that sit on a chair. We aim to understand how these oscillatory episodes emerge online as a function of variations in specific navigational demands during the trial such as familiarity with the path, location within the path, distance to the next turn, and accuracy.

## METHODS

### Subjects

The experiment was run in accordance with the Declaration of Helsinki, under approval of the CEIC Parc de Salut Mar (Universitat Pompeu Fabra) institutional review board (approval number 2023/10922/I). All participants gave prior written informed consent, were naïve to the hypothesis of the experiment, and were compensated with 10 € per hour (in 30min fractions). Participant exclusion criteria were: history of epilepsy, being under medication for neurological or psychiatric disorders, and susceptibility of suffering motion sickness, screened online using the Cybersickness Susceptibility Questionnaire (Freiwald et al., 2020). 24 subjects were recruited for the experiment (13 female, 25±5 years), and all were included in the final sample. The experimental session had an approximate duration of 3 hours.

### Virtual Reality (VR) environment

The VR environment was programmed using the Unity Real-Time Development Platform (Unity Technologies Inc., version 2020.3.19f1,) and was presented to subjects via a Head Mounted Display HTC Vive (HTC corporation) (Figure 1C). During navigation, subjects were seated and drove a car through streets using a steering wheel (Figure 1C). and one foot-pedal Logitech G29 (Logitech International S.A.)

**Figure 1:**
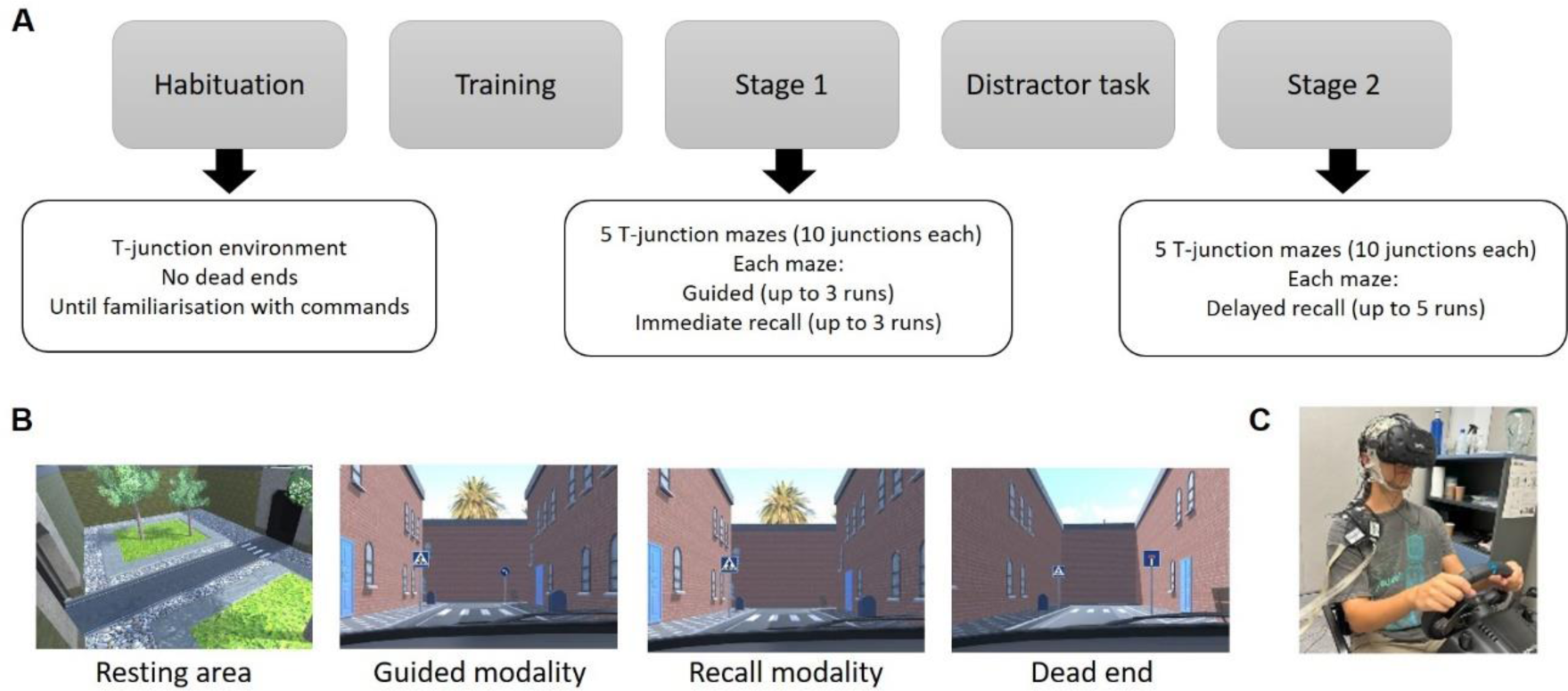
**A:** Experimental procedure: Subjects entered first a habituation environment consisting of T-junctions, they were instructed on using the steering wheel and pedal. They were allowed to stay in the environment until they were familiar with the commands. Next, participants performed a training, consisting of a short version of the mazes presented in Part I. In Part 1, 5 T-Junction mazes (10 junctions each) were presented in randomised order. Each of the mazes was presented consecutively in 2 modalities: Guided and Immediate recall. The order of the maze presentation was randomised across subjects. For each modality, subjects had up to three attempts to solve the maze. After completion of Part I, subjects took a break, removed the HMD and performed a distractor task: they played a Timeline card game (Zygomatic) against the experimenter. Finally, subjects entered the VR again and performed the Delayed recall: they were presented with the mazes learnt in Part I, in randomised order. Each maze was presented without directional cues and subjects had up to five attempts to solve it. During all of the virtual navigation, white noise was presented to the subjects by means of earbuds. B: Snapshots of the VR environment: Resting area that subjects entered after an error or after successfully solving a maze. Subjects were allowed to rest there as long as they required before entering the next maze. In the Guided modality, directional cues (traffic signs with arrows) indicated which was the correct side to turn. In the Immediate and Delayed recall modality, the mazes were the same as the ones presented in the Guided modality, but directional cues were removed. When subjects committed an error, they entered a dead end and were teleported to the resting area. **C**: Image of a subject performing the task.

The VR environment included three scenarios: resting area, habituation environment, and T-Junction mazes. Subjects entered the resting area (Figure 1B) on initiation of the experiment, where they could adjust their position, and align the virtual dashboard and the steering wheel with their own posture. In the resting area, a traffic light turned from red to green to indicate that subjects were allowed to proceed to the next scenario at a time of their choosing. Next, the habituation environment consisted of navigating a sequence of T-Junctions. Each arm of the maze consisted of a straight road ending with a T-crossing. Participants pressed the foot pedal to move forward and were required to turn to left or right in time, before reaching a front wall, at the crossing. In this scenario subjects could freely choose to turn left or right. The onset of the turn could happen at any moment after crossing and the crossway corresponded to the moment in which subjects started to rotate the steering wheel to indicate the selected turn. The purpose of the habituation environment was to familiarise subjects with the VR scenarios and the function of the steering wheel and pedal. After each turn, another T-junction started without a break in the visual flow, and subjects were allowed to drive in the habituation environment for as long as they required.

Finally, the T-Junction mazes environment used in the experiment, consisted of paths of 10 consecutive T-Junctions, similar to the habituation scenario. However, for each different maze there was a unique combination of left (L) and right (R) turns (for instance: LLRLRRRLRL) to reach the end point (target destination). If subjects took a wrong turn in any of the modalities (Guided, Immediate, or Delayed recall), they entered a dead end (Figure 1B) and were teleported to the resting area. Otherwise, they proceeded to the next arm of the maze until the end point, which was signalled by a tunnel. Once they successfully reached the end of a maze, subjects were teleported to the resting area. We created five different mazes for the experiment, plus an additional training maze (consisting of 4 T-junctions). Each of the mazes simulated a succession of city-like streets with a unique, distinctive, combination of the following features: wall colour and texture, pavement texture, street elements (benches, lamps, plants), and window and door styles, all uninformative as to the correct path (Figures 1B-D).

### Experimental protocol

The experiment took place in a laboratory room (CBC labs, Universitat Pompeu Fabra, Barcelona, Spain). On arrival, subjects were screened for susceptibility to cybersickness adverse symptoms (headache, blurred vision, eye strain) using the Simulator Sickness Questionnaire (Kennedy et al., 1993). None of the subjects reported severe symptoms and, therefore, all proceeded to the experiment. Next, subjects were seated on a chair, fitted with the VR goggles, and the SN paradigm started (Figure 1C). The paradigm was divided in two parts, separated by a distractor task. In Part I, the five mazes were learnt. For each maze, subjects first navigated the entire maze with direction cues indicating the correct turns (Guided mazes). Subjects had up to threeattempts to correctly follow the path, but most participants completed the maze at the first attempt in this stage. Immediately after, they had three attempts to solve the same maze without directional cues (Immediate recall mazes). Each attempt ended either upon an error or after successful completion. After studying the five different mazes, participants were allowed to exit the VR environment to take a resting break, during which they were invited to play the memory game Timeline (distractor task) for 15 minutes. After the break, Part II took place back in the VR environment. Subjects had to solve once again the five mazes learnt in Part I, now presented in random order and without directional cues (Delayed recall mazes). EEG was recorded during both parts of the experiment, including five minutes recordings of resting-state activity (eyes closed) prior to the task.

Upon finishing, subjects filled out the Simulator Sickness Questionnaire again (Kennedy et al., 1993) to control for adverse symptoms due to the VR, and an immersion questionnaire adapted from Cárdenas-Delgado et al. (2017).

### EEG acquisition and pre-processing

EEG was recorded during Part I and Part II of the experiment at a sampling rate of 500 Hz using a BrainAmp amplifier and BrainRecorder software (BrainProducts, GmbH, Munich, Germany) with 64 active electrodes (actiCAP, BrainProducts, GmbH, Munich, Germany) placed according to the 10-10 system. The ground electrode was placed at the left mastoid and the online reference on the tip of the nose. The TP9 electrode was moved to the Ground position in the cap, TP10 to the right Mastoid, and Fp2 to the Reference position in the cap. Two electrodes were used for horizontal and vertical electrooculogram, specifically FT10 was used as the VEOG (right) and FT9 as the HEOG (left). Impedances were kept below 10 kΩ.

Pre-processing and segmentation were run using EEGLAB (Delorme & Makeig, 2004), Fieldtrip (Oostenveld et al., 2011) and custom-written code. First, noisy channels were detected using a combination of visual inspection and automatic procedures implemented with EEGLAB plugin *clean_rawdata* (flat channel for more than 10 seconds, high frequency noise above 4 standard deviations, correlation with neighbouring channels below 0.8). Noisy channels were discarded. Second, Independent Component Analysis (ICA) (Artoni et al., 2018) (extended *runica*) was used on continuous data to correct visual artefacts. *ICLabel* EEGLAB plugin (Pion-Tonachini et al., 2019) was used for the automatic labelling and discarding of components corresponding to blinks and eye movements, as well as muscle artefacts (cut-off threshold p> 0.975). Third, the data was re-referenced to the right mastoid and a high pass filter (Butterworth, order 4, non-causal, cut-off frequency 0.1 Hz) and a low pass filter (Butterworth, order 4, non-causal, cut-off frequency 80 Hz) were applied. Finally, pre-processed data was segmented in trials from -6500 to 500 ms relative to the turn onset, for the Immediate and Delayed recall mazes. The rejected channels were reconstructed using the data of neighbouring channels and spherical splines. Data recordings at rest (eyes closed) were pre-processed in the same way as the spatial navigation task data and subsequently divided in epochs with the same duration of the task epochs. Across subjects, 4±2 electrodes were marked as bad and interpolated 2±1 visual components, and 3±2 muscle components were detected and discarded.

### EEG analysis

EEG analysis was run using Fieldtrip (Oostenveld et al., 2011) and custom-written code. Data was downsampled to 250 Hz. The frequencies of interest were theta (3 to 8 Hz) and alpha (8.5 to 14 Hz; range chosen to avoid spectral overlap with theta band, the main frequency of interest). We used the *fBOSC* toolbox (Seymour et al., 2022) to identify oscillatory episodes.

Power was extracted in a time-resolved fashion within each epoch for frequencies in the 2 to 40 Hz range, 0.5 Hz steps, by means of Morlet wavelets (6 cycles). Oscillatory episodes were defined as segments of signal where power in the specified frequency exceeded the 99^th^ percentile of background activity for at least three cycles. For each trial, electrode, and subject, we calculated the abundance (Seymour et al., 2022) of oscillatory episodes as the percentage of total time in which the signal presented oscillatory episodes in a specified band (abundance is virtually equivalent to *p-episode*, in Caplan et al. 2001). We then detected the electrodes that presented higher theta and alpha abundance at the group level during spatial navigation. These electrodes were then selected for further single-subject and single-trial analyses.

### Statistical analysis

We used linear mixed models (LMMs) for the statistical analysis, so that individual variability can be modelled in a straightforward manner using random factors. Because LMMs can handle missing values and are robust to unbalanced designs they are convenient to simultaneously consider all the factors that can have an impact on the structure of the data (Verbeke & Molenberghs, 2009).

We divided the trials recorded during the spatial navigation task in segments of two seconds centred at times -5000 ms to -1000 ms before the turn (t=0) in steps of two seconds. Abundance, defined as the percentage of time in which an oscillatory episode was detected in the segment, was our variable of interest. Abundance is a continuous variable that can range from 0% (no oscillatory episode detected in the segment) to 100% (the whole segment consisted of oscillatory activity). Note that we excluded from the analysis the intervals -6500 to -6000 ms and 0 to 500 ms. Those intervals were excluded because, on one hand, we wanted to consider only data previous to the turn and, on the other hand, the abundance at the edges of the EEG segment can be underestimated^2^.

We used a model with abundance as the dependent variable, participant as the random factor, and the fixed factors described in Table 1. The factors selected contain information about different aspects of the task. The ability to correctly recall learnt mazes is described by factors Correct, that alludes to the correct or incorrect recall of the specific junction and Solved, that describes the performance for the whole maze, whether the maze was solved or not. The other fixed factors (Stage, Run, Junction and Time) describe the characteristics of the task at different time scales. Stage refers to a very long time scale (approximately one hour): whether the trial is part of an Immediate Recall maze, in the first part of the experiment, recalling the paths that were just learnt or part of a Delayed Recall maze, in the second part of the experiment, and has to recall the path of one of the five mazes learnt in the first part of the experiment. Factor Run indicates how many consecutive attempts to solve a maze the subject performed, two runs of the same maze would be separated by a time ranging from 10 to 100 seconds. Factor Junction encodes the position in the maze, whether the subject is starting the maze (Junction 1) or close to the end of the maze (Junction 10). Two consecutive Junctions were separated by approximately 10 seconds. Finally, factor Time indicates, in a specific junction how far from the decision point is the subject.

**Table 1:** Factors included in the LMM.

| Factor | Range | Description |
| --- | --- | --- |
| Stage | 1 or 2 | 1: Immediate recall trials, 2: Delayed recall trials |
| Run | 1,2,3 (Immediate recall)<br>or 1,2...5 (Delayed recall) | Attempt number to solve a given maze at a given stage. |
| Junction | 1, 2...10 | Turn number that the subject is about to take on a given maze. Corresponds to the location within a path. |
| Correct | 0 or 1 | If true (1), subject correctly recalled the direction to turn in the upcoming junction |
| Solved | 0 or 1 | If true (1), the considered epoch is part of a run in which the subject provided 10 consecutive correct decisions. |
| **Time** | -5000, -3000, -1000 | Time (in ms) before the junction and decision of the turn direction, related to the distance to the upcoming turn. |

Statistical analysis was performed in R (R Core Team, 2022) with functions from the package *glmmTMB* (Brooks et al., 2017), using ordered beta family for the fit (see Supplementary Materials and Methods). In order to select the model that best described the data, we started from the full model:

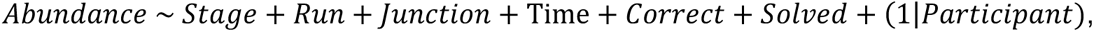

and used the dredge function to build all possible models with this set of factors and *MuMIn* R package (Bartoń, K., 2009) to rank them according to second order Akaike Information Criterion (AIC) (Burnham et al., 2010). Significance of the factors was assessed by means of ANOVA (type III *Χ*^2^ test). Packages *superheat*(Barter & Yu, 2018) and *ggplot2*(Hadley Wickham, 2016) were used for visualization.

## RESULTS

### Behavioural results

All subjects reached the end of all Guided mazes. Most subjects (N=20) required only one attempt in all five mazes, three subjects committed an error in one out of the five mazes, and one subject committed an error in two runs of one of the five mazes. Subjects correctly recalled 8.31±1.15 junctions in the Immediate recall mazes, and 5.95±1.71 in the Delayed recall mazes. In both cases, the number of consecutive correct junctions increased with each run (Figure 2).

**Figure 2:**
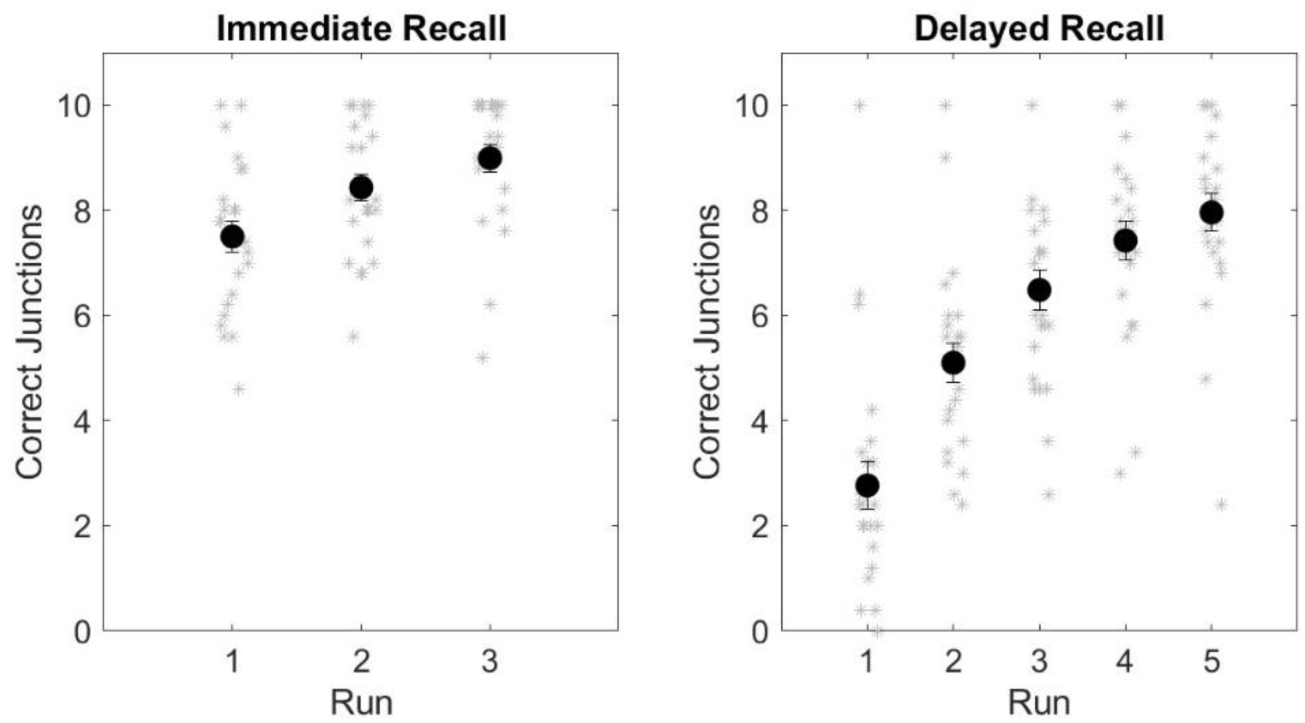
Mean number of correct junctions recalled in the Immediate recall mazes (left) and Delayed recall mazes (right). Grey dots correspond to individual mean performance and black dots correspond to the mean across subjects, error bars correspond to the standard error of the mean.

### Oscillatory episodes

We used an oscillation detection method to estimate the abundance of oscillatory episodes in the relevant frequencies and scalp locations (see Methods for details on the analytical pipeline and statistical approach). At the group level, most of the oscillatory episodes (Figure 3F) were observed at frequencies below 15 Hz (Figure 3A). The abundance of theta (3 to 8 Hz) episodes during the spatial navigation task was higher in frontal channels, with the maximum at electrode Fz (Figure 3B). Both the power spectrum of the channel and abundance of oscillatory episodes displayed a double peak, with the maximum in the theta range, at 6.5 Hz, and a second peak in the alpha range, at 9 Hz (Figures 2D and Supplementary Figure S2A). Mean theta abundance was 20±12%, and the episodes had a mean duration of 5±1 cycles (880±140 ms). By contrast with task-related activity, the power spectrum and abundance of channel Fz at rest presented a single peak at 9 Hz (Figure 3E and Supplementary Figure S2B). The abundance of theta episodes in channel Fz at rest was 14±9%, significantly lower than during Immediate and Delayed recall task trials (two-tailed paired t-test, alpha level 0.05, *t*(23)= -2.92, *p*=.008, Cohen’s d=-0.6).

**Figure 3:**
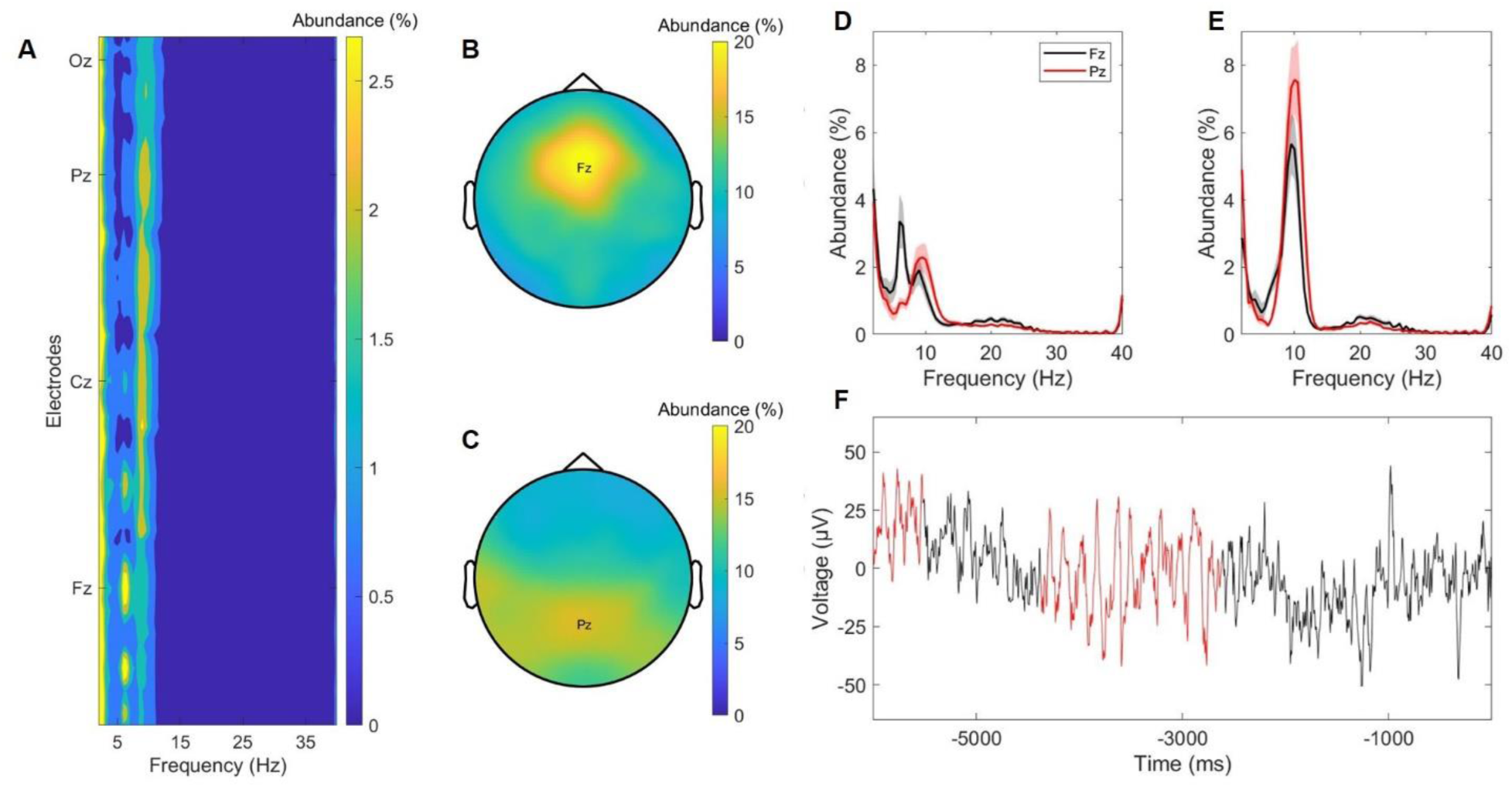
**A:** Distribution of episodes across frequency and electrodes. **B:** Topographical distribution of theta (3 to 8 Hz) episodes. **C:** Topographical distribution of alpha (8.5 to 14 Hz) episodes. **D:** Episode abundance for channels Fz (black) and Pz (red) during the spatial navigation task. **E:** Episode abundance for channels Fz (black) and Pz (red) at rest. **F:** Example of the EEG trace during the spatial navigation task for a subject (black) with two oscillatory episodes identified (red). Time 0 corresponds to onset of the decision.

Alpha (8.5 to 14 Hz) episodes were more abundant over parietal areas, with the maximum at electrode Pz (Figure 3C). The power spectrum and the abundance of oscillatory episodes displayed a single peak in the alpha range, at 9 Hz (Figure 3D and Supplementary Figure S2A). Mean alpha abundance was 15±10%, alpha mean episode duration was 5±1 cycles (480±90 ms). At rest, both the power spectrum and alpha abundance at channel Pz presented a clear single peak at 9 Hz (Figure 3E and Supplementary Figure S2B). The abundance of alpha episodes during rest (46±20%) was significantly higher than during Immediate and Delayed recall task trials (two-tailed paired t-test, alpha level 0.05, *t*(23)= 6.34, p<.001, Cohen’s d=1.3).

The abundance of oscillatory episodes was highly variable between subjects for both theta and alpha bands (Figure 4). In order to determine whether oscillatory activity (irrespective of frequency band) was easier to detect in some subjects, we calculated the correlation between theta and alpha abundance across subjects. There was no correlation between theta and alpha abundance across subjects (*r*=.02, *p* =.9).

**Figure 4:**
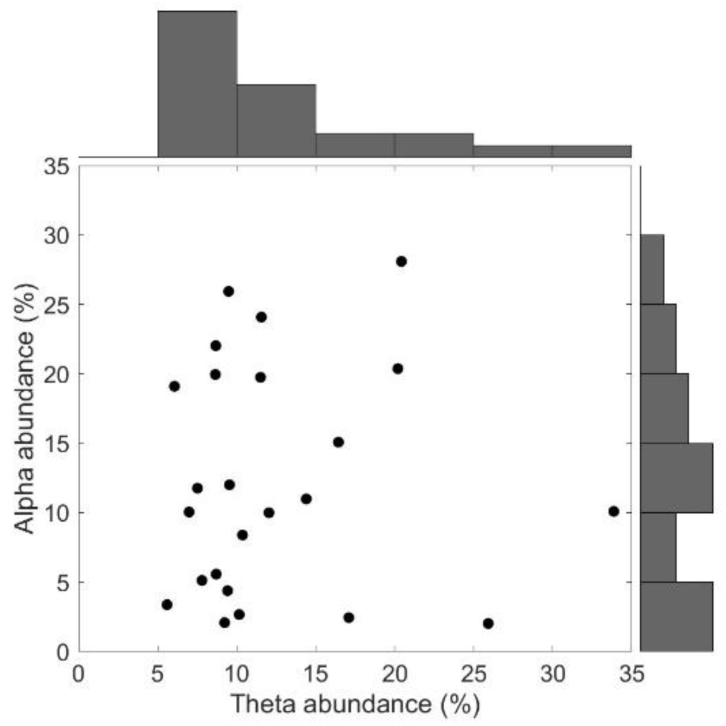
Individual abundance values averaged across time, electrodes and trials for theta and alpha bands. Each point corresponds to one subject. Distribution of oscillatory episodes is shown in the top (theta) and right (alpha) histograms.

Theta episodes were more abundant in Delayed recall trials, compared to Immediate recall trials (Figure 5A-E). Episode abundance increased with run number (Figure 5A), and it was larger in the central junctions of a given maze, especially at early times within the approach to the junction (Figure 5B, Figure 5C, and Figure 6). Correct decisions resulted in decreased theta episode abundance, only for Immediate recall mazes (Figure 5D). Finally, theta abundance did not differ between solved and unsolved mazes (Figure 5E).

**Figure 5:**
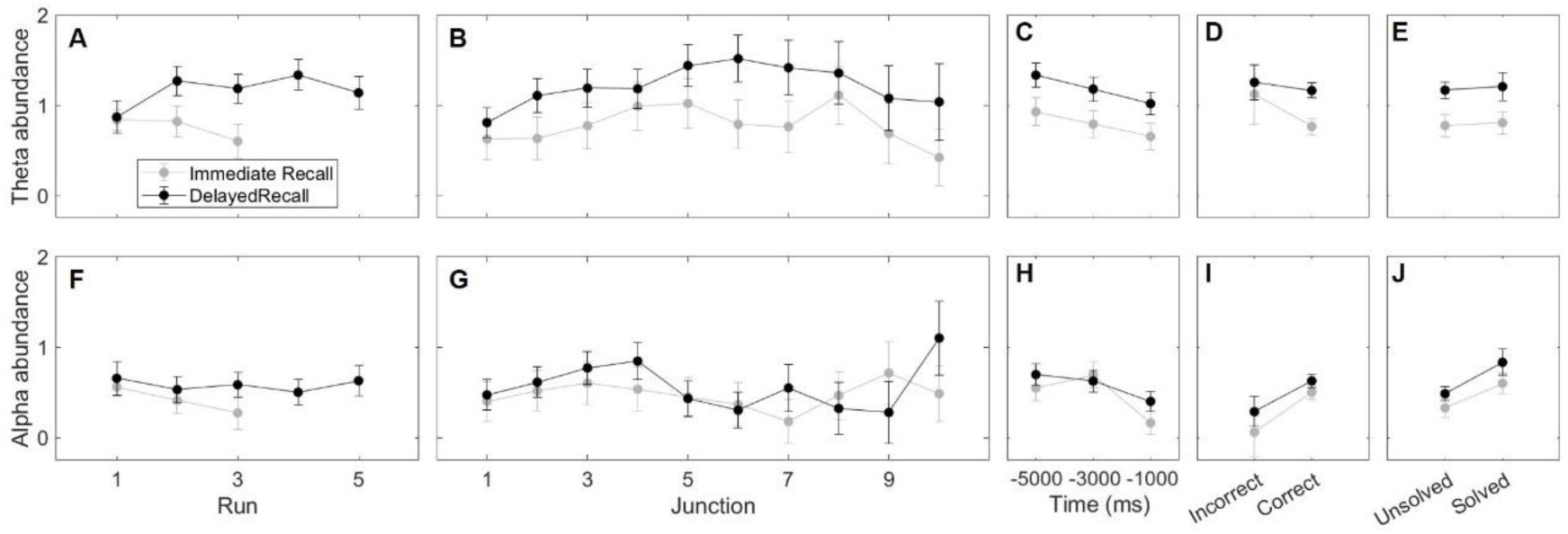
Mean episode abundance for Immediate recall (grey) and Delayed recall (black) mazes. Values are z-scored with respect to the values shown in Figure 3. Theta band **A** and **F:** Abundance as a function of Run. **B** and **G:** Abundance as a function of Junction within a maze. **C** and **H:** Abundance as a function of Time within the approach to a junction turn, averaged in 2 s bins centred at -1, -3 and -.5 s prior to the turn. **D** and **I:** Abundance as a function of decision accuracy in the upcoming turn. **E** and **J:** Abundance averaged across the whole maze, as a function of whether the maze was solved correctly or not. Error bars indicate the 95% CI.

**Figure 6:**
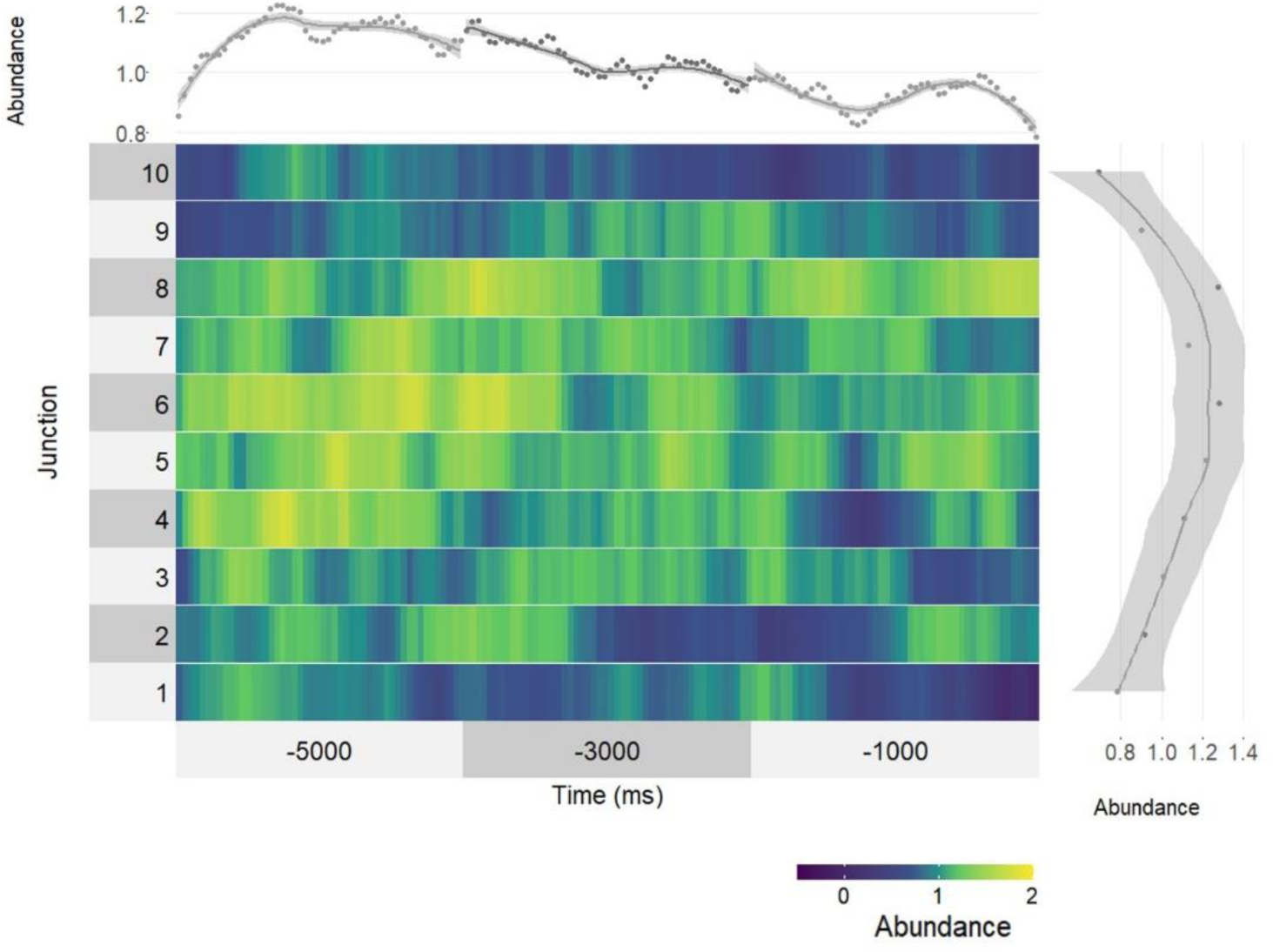
The central plot displays the time-resolved mean theta episode abundance values as a function of Junction within a maze (y-axis) and Time to decision within a Junction approach (x-axis). For each subject, Abundance was z-scored with respect to the individual theta episode abundance values displayed in Figure 3. The top plot displays abundance as a function of Time to decision within a junction approach averaged across all junctions, and the right margin plot displays abundance as a function of Junction, averaged across Time within the Junction approach. Lines correspond to locally estimated scatterplot smoothing and shaded areas to the standard error of the estimation.

Contrary to theta episodes, the abundance of alpha episodes did not seem to strongly differ between the Immediate and Delayed recall stages (Figures 5F-J and 7). Alpha episodes increased monotonically during the first four runs (Figure 5F) and were more abundant in the first junctions of the mazes, especially at the early time segments within the junction (Figure 5G, Figure 5H and Figure 7). Successful navigational decisions were preceded by increased alpha abundance at junction level (correct compared to incorrect decisions, Figure 5I) as well as at maze level (solved compared to unsolved mazes, Figure 5J).

**Figure 7:**
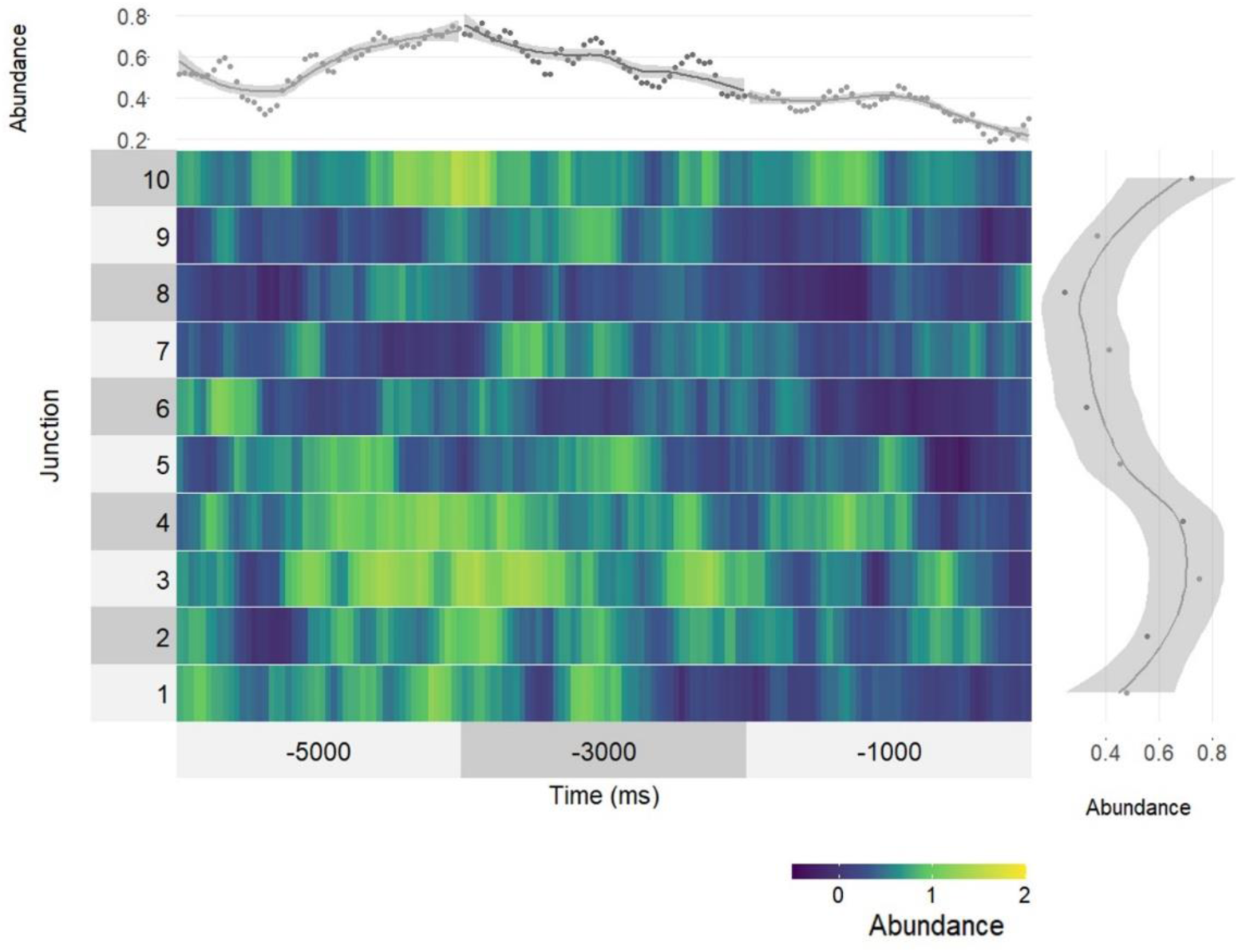
The central plot displays the time-resolved mean alpha episode abundance values as a function of Junction within a maze (y-axis) and Time to decision within a Junction approach (x-axis). For each subject, Abundance was z-scored with respect to the individual alpha episode abundance values displayed in Figure 3. The top plot displays abundance as a function of Time to decision within a junction approach averaged across all junctions, and the right margin plot displays abundance as a function of Junction, averaged across Time within the Junction approach. Lines correspond to locally estimated scatterplot smoothing and shaded areas to the standard error of the estimation.

### Statistical analysis

For the theta band, electrode Fz, the model that best described the data was (see Supplementary Materials, 2.2. Model selection):

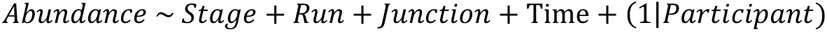

Stage, Junction and Time had a significant impact on theta Abundance (Table 2), whereas Run, Correct response and Solved did not have an impact on theta Abundance. Post-hoc tests (Supplementary Table 3) indicated that Abundance was significantly higher in the Delayed recall mazes compared to Immediate recall mazes, in the middle of the maze (Junctions 5 and 6) compared to the beginning or end of the maze (Junctions 1 and 10), and at the early times of the trial compared to medium and late times. Interestingly, these effects were mostly driven by Delayed recall trials (Supplementary materials and methods: 3.3. Models for Immediate recall and Delayed recall trials) whereas in Immediate recall trials only the response (Correct or Incorrect) had an impact on oscillatory theta episodes. Predicted Abundance values for the significant factors included in the model selected for theta band are shown in Figure 8.

**Figure 8:**
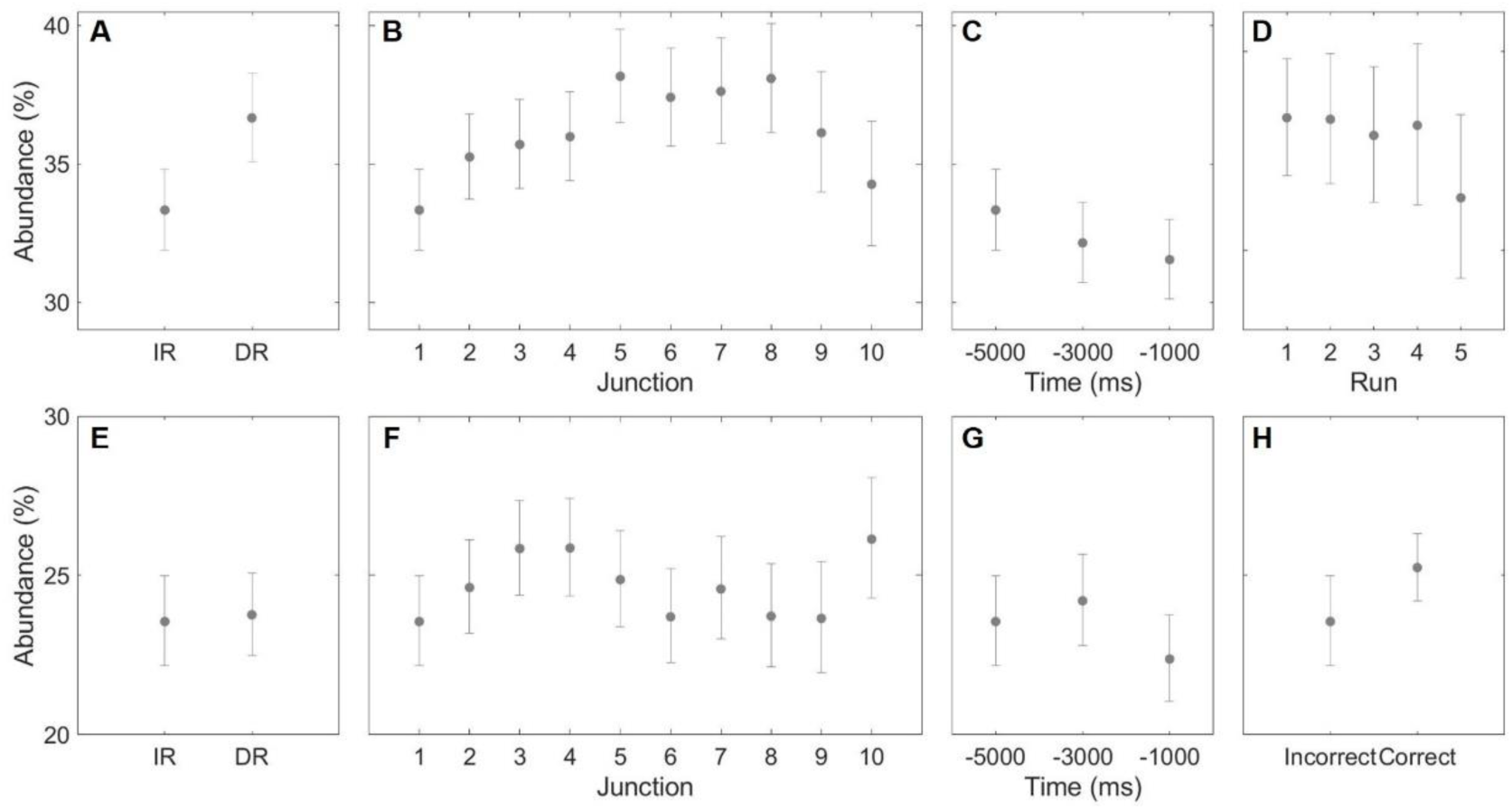
Predicted abundance values for theta (**A** to **D**) and alpha (**E** to **H**) frequency bands according to the linear mixed models selected for each frequency band (see Table 2 (theta) and Table 3 (alpha)). **A** and **E**: Immediate (IR) and Delayed (DR) recall trials. **B** and **F**: As a function of junction. **C** and **G**: As a function of time to decision. **D**: As a function of run. **H**: For correct and incorrect decisions.

**Table 2:** ANOVA (type III, Wald) for the model describing Abundance in the theta band.

| | $\chi^2$ | d.f. | *p* |
| --- | --- | --- | --- |
| **Participant** | 59.62 | 1 | <.001 |
| **Stage** | 23.68 | 1 | <.001 |
| **Run** | 8.41 | 4 | .08 |
| **Junction** | 45.3 | 9 | <.001 |
| **Time** | 16.15 | 2 | <.001 |

**Table 3:** ANOVA (type III, Wald) for the model describing Abundance in the alpha band.

| | $\chi^2$ | d.f. | *P* |
| --- | --- | --- | --- |
| **Participant** | 110.88 | 1 | <.001 |
| **Stage** | 8.99 | 1 | .003 |
| **Junction** | 34.82 | 9 | <.001 |
| **Time** | 28.73 | 2 | <.001 |
| **Correct** | 18.43 | 1 | <.001 |

For the alpha band at electrode Pz, the model that best described the data was (see Supplementary Materials, 2.2 Model selection):

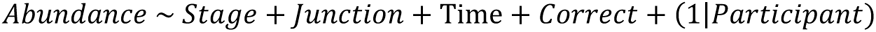

Abundance in the alpha range was modulated by Stage, Junction, Time, and Turn accuracy (Correct) (Supplementary Table S4), whereas Run and Solving the maze had no impact on alpha abundance. Post-hoc tests (Supplementary Table S9) revealed an increase in alpha episodes during Delayed recall mazes compared to Immediate recall mazes and in the middle Junctions of the maze (Junctions 3 and 4) as compared to the first Junction. We also observed a decrease in abundance of alpha oscillatory episodes as the subject approached the decision point within a trial. Finally, alpha abundance was higher in trials where subjects made a Correct turn decision, compared to Incorrect decision trials. When we separately modelled Immediate and Delayed recall trials, the pattern of results for Delayed recall trials was similar to the one observed for all trials, whereas Time was the only factor significant for Immediate recall trials (Supplementary materials and Methods: 3.3 Models for Immediate recall and Delayed recall trials.). Predicted Abundance values for the significant factors included in the model selected for alpha band are shown in Figure 8.

Summarizing, both frontal theta and parietal alpha oscillatory episode abundance increased during Delayed recall as compared to Immediate maze recall. In addition, although for both frequency bands abundance of oscillatory episodes increased as subjects advanced through the sequence of junctions along a maze, alpha abundance peaked before theta abundance (Figure 9), and only theta abundance decreased significantly by the end of the maze (Figure 9). In addition, parietal alpha oscillatory episodes abundance differed as a function of upcoming turn accuracy, with abundance being higher before correct as compared to incorrect turns (Figure 9). Finally, neither the model for theta nor alpha bands revealed significant interactions between the factors included in the selected models (see Supplementary Information 4.4 Interactions).

**Figure 9:**
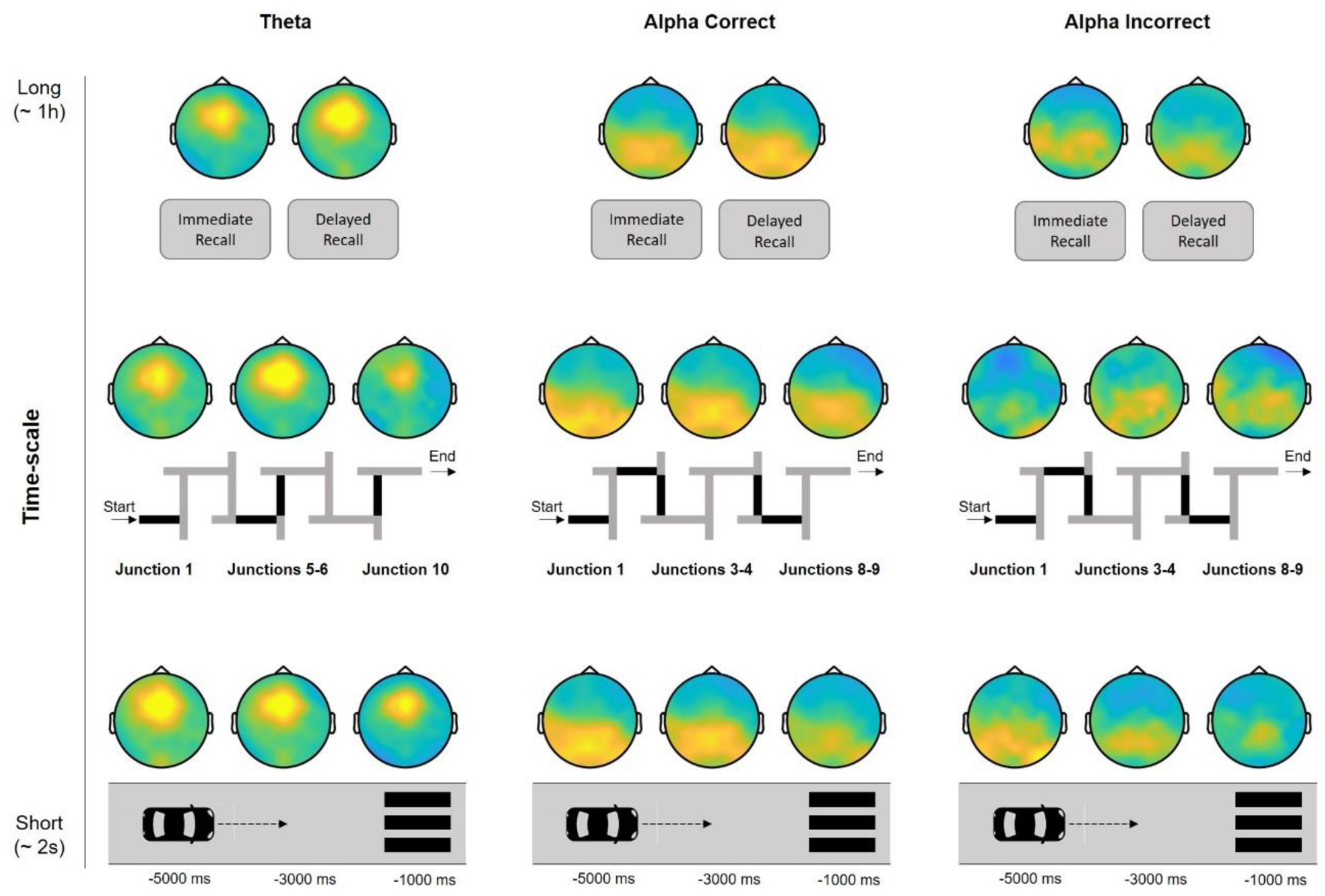
Summary of episode abundance across different time scales. At a long time scale, both Frontal theta and Parietal alpha episode abundance increased on Delayed Recall trials compared to Immediate recall trials. Within a maze time scale (approximately 2 minutes), frontal theta episodes’ abundance significantly increased in the middle part of the maze (in particular, at junctions 5 and 6) compared to the start and end of the maze, whereas Parietal alpha increased on junctions 3 and 4 as compared to the first junction. In a junction, episode abundance for both Frontal theta and Parietal alpha was maximal at the start of the junction and progressively decreased when subjects approached the decision point. Lastly, alpha episodes’ abundance was significantly higher for trials preceding a correct decision compared to an erroneous one.

## DISCUSSION

The present results bring to light the relevance of moment-to-moment episodes of oscillatory neural activity during a spatial memory task in humans, at a single-trial level. The relevance of individual bouts of theta activity has been widely assumed in the literature on human spatial navigation, but rarely addressed, especially using non-invasive methods. Interestingly, we observed a clear spatial differentiation between the loci of the most abundant theta and alpha episodes: Theta abundance peaked at frontal locations, whereas alpha abundance was maximal at parietal sites. The occurrence episodes of frontal theta and parietal alpha oscillatory activity was found to be tightly related to relevant events during navigation. We observed differences in the dynamics of episode abundance both at long (along the progression of junctions across the maze) and short time scales (along the path toward a junction within each arm of the maze). We observed increases in alpha episodes at early junctions along the maze, whereas theta abundance was highest in the middle junctions. Within a single junction arm, theta episodes were most abundant early on and decreased monotonically as subjects approached the turning point, whereas alpha episodes were sustained throughout the arm’s length and decreased only close to the turn. Finally, only alpha activity correlated with successful navigational decisions, specificallyan increased abundance of alpha oscillatory episodes anticipated correct turns.

More generally, our findings reinforce current views that spatial navigation is supported by an extended network of brain areas. Although probably biased by the vast and important literature using electrophysiology in rodent models, traditional theories consider the hippocampus to be the key structure in spatial navigation (O’Keefe & Nadel, 1978). This view is, at least in part, currently being challenged by newer theories. First, these new theories tend to attribute a more general role to the hippocampus, which would serve to create cognitive maps, including but not restricted to, spatial navigation (Buzsáki & Moser, 2013). Second, although reserving a prominent role to hippocampal activity, new findings have revealed a wide network of additional brain areas involved in the computations necessary for spatial navigation (Ekstrom et al., 2017). This latter view is supported by neuroimaging studies indicating that oscillatory activity at different frequency bands (Caplan et al., 2001; Chrastil et al., 2022; Jacobs et al., 2010; White et al., 2012; Zhu et al., 2022) and locations (Baumann & Mattingley, 2021; Caplan et al., 2001; Epstein et al., 2017) is involved in spatial navigation, including our study, in which both frontal theta and parietal alpha varied as a function of different aspects of the spatial navigation task.

In our study, frontal theta abundance was related to global properties of the navigation task. At the longest time scale tested, we observed increased abundance of theta episodes during Delayed Recall as compared to Immediate Recall, in agreement with the seminal findings of Kahana et al. (1999) using intra-cranial recordings in humans. This increase of theta episodes could be explained by an anticipation of the properties of the task: Theta episodes are more abundant in the more difficult Delayed Recall trials, possibly revealing the effortful retrieval of previously encoded paths. At a shorter time scale, involving the junction-to-junction sequence within a maze, theta abundance was observed to increase precisely in the middle junctions. A feasible interpretation of this result aligns with the classical serial position effect in memory for sequences, a phenomenon that has also been reported to apply in the context of spatial navigation (Hilton et al., 2023). In short, T-Junction mazes can be conceptualised as an ordered sequence of binary decisions (for instance LRRLRLLLRR). According to the serial position effect, there would be a facilitation in the encoding and subsequent recall of first and last items of the sequence (due to primacy and recency effects respectively) compared to the middle parts of the sequences. Therefore, in our experiment, increased theta abundance in the middle part of the mazes can be linked to the increased difficulty of retrieval of these middle junctions (Caplan et al., 2001; Kahana et al., 1999). At the shortest time scale, as the participants progressed toward a junction turn, we observed an initial heightened abundance of theta episodes followed by a monotonic decrease. This might suggest that subjects accessed encoded path information at early moments in the junction and, once a navigational decision was made, encoded information was not further accessed. For theta episodes, the results indicate a robust pattern that associates more abundant and stronger theta oscillatory episodic activity with access to spatial information during navigation (see Supplementary Information 5.1 Models for frontal theta power and episode power).

Although previous non-invasive EEG studies have observed a positive correlation between theta band activity and performance in spatial navigation tasks (Bauer et al., 2021; Chrastil et al., 2022; White et al., 2012), we found that abundance of theta episodes was not significantly different prior to correct and incorrect navigational decisions, in line with previous studies (Caplan et al., 2001). A possible explanation for the discrepancy could be differences in the spatial navigation task: In the studies finding a positive correlation between theta and spatial navigation performance the navigational choices for the subjects were less limited than in our study, where subjects had to choose between only two options. Therefore, the sample of correct responses in our study probably contains trials in which subjects chose the correct decision by chance, possibly blurring the comparison between correct and incorrect choices.

Contrary to theta, the abundance of alpha episodes correlated with successful navigational decisions: increased parietal alpha abundance was found in trials preceding a correct decision, a result that aligns with a recent study linking improved performance in spatial navigation with increases in parietal alpha activity (Chrastil et al., 2022). At present, alpha activity is considered to reflect active inhibition of task-irrelevant representations, such as distractor information (Klimesch, 2012): consistent findings from working memory studies using sequences of simple stimuli (such as letters and numerals) indicate an increase of alpha power during the retention interval especially when distractor information must be suppressed, and a positive correlation between alpha power and working memory load (Jensen, 2024). Following these results, alpha activity has been proposed to protect memory maintenance against distracting information (Bonnefond & Jensen, 2012). The pattern of alpha episodes in our study would be compatible with this protective role. In line with this idea, increased posterior alpha oscillations have been suggested to signal switches between internal and external attention (Benedek et al., 2014; Magosso et al., 2021). Interestingly, this has also been confirmed in VR immersive tasks (Magosso et al., 2019). This interpretation aligns well with our results: considering the absence of relevant landmarks inside the mazes, participants had to rely heavily on episodic memory traces. Therefore, increased alpha abundance on correct decision trials could indicate trials in which subjects managed to focus on internal attention more successfully. Increases in alpha episode abundance in Delayed Recall compared to Immediate Recall trials could be explained similarly: In the second stage of the experiment, accessing internal representations of the previously learnt paths was the optimal strategy to solve old mazes. Therefore, an increase in alpha in sensory areas would indicate suppression of incoming information possibly distracting from access to internal representations. This suppression would be reflected in more abundant and stronger parietal alpha episodes (see Supplementary Information: 5.2. Models for parietal alpha power and episode power).

It could be argued that the design of our experiment conflates Immediate vs. Delayed recall conditions with total time on task, complicating the interpretation of variations in theta and alpha abundance across experimental stages. Increases in overall alpha and theta with time on task have previously been associated with fatigue, drowsiness, and task detachment (see Tran et al. (2020) for a systematic review). However, when discussing time-in-task effects, it is important to dissociate task-related theta from task-unrelated theta activity. Arnau et al. (2021) reported that whilst inter-trial theta activity increased with time on task, task-related activity was reduced; therefore, fatigue and detachment from the task is in fact associated with reduced theta, an interpretation that aligns with models relating theta oscillations to cognitive control (Cavanagh & Frank, 2014). Moreover, it is important to note that, in the present experiment, performance did not decrease with time on task (see Supplementary Materials), ruling out a time-dependent task disengagement effect. Therefore, although we cannot completely rule out a contribution of time-on-task, evidence suggests that the increased difficulty (and higher demand on memory) is the most likely explanation for the increase of theta abundance on Delayed recall conditions.

One important aspect of the current study is that we used estimations of purely oscillatory activity from the EEG signal, and therefore the abundance data is not contaminated by changes in wide-band activity. It is now accepted that brain oscillatory activity emerges in short intermittent episodes distributed stochastically in time, and with variable duration in cycles, superimposed on the background wide-band aperiodic activity (S. R. Jones, 2016; Seymour et al., 2022). Targeting these oscillatory episodes in a time-resolved fashion (e.g., at a single-trial level) is therefore essential to capture the relationship between neural oscillations and behaviour. Yet, despite its paramount importance, the distinction between oscillatory and aperiodic activity is often neglected (Gerster et al., 2022). Although prevalent theories relating theta to spatial navigation assume that the theta activity involved in information organization and transmission is an oscillation (Buzsáki, 2005; Ekstrom et al., 2017), many electrophysiological studies of spatial navigation do not isolate purely oscillatory activity from aperiodic activity, making it impossible to discriminate changes related to oscillatory activity from changes related to broadband activity, that have been shown to vary with the task (Gilden, 2001; Ouyang et al., 2020; Podvalny et al., 2015; Waschke et al., 2021). Our study is in line with the few empirical studies that, to our knowledge, have associated purely theta oscillatory activity with spatial navigation in humans (Caplan et al., 2001; Du et al., 2023; Gehrke & Gramann, 2021; Kahana et al., 1999).

What is more, we used an exploratory approach to inspect the data at a single-trial level. This allowed us to capitalise on the complexity of the data at different levels of granularity. Dynamics of oscillatory episodes were rich, presenting patterns at different time scales that can be related to the difficulty of the task (theta band), access to encoded information (theta band), and switching and focusing attention (alpha band). On the one hand, we found that theta correlates with global demands of the task, as increased theta episode abundance was observed in the more difficult parts of the task at various time scales. At a longer time scale it increased from Immediate to Delayed recall; at a medium time scale theta abundance increased in the middle of the path compared to the beginning and end; and, at a short time scale within a single junction, frontal theta abundance signalled early access to encoded information in anticipation of the impending navigational decision. On the other hand, alpha oscillatory activity can generally be interpreted as indicating switches to/from internal attention, with increases in parietal episodes possibly signalling the inhibition of distracting sensory information to facilitate selective access to internal episodic representations.

In conclusion, the results of our study provide crucial support for long-standing theories linkingtheta oscillatory activity with access to stored information relevant for spatial navigation. We addressed brain oscillations during task-relevant events at moment-to-moment single-trial level, and revealed rich temporal, spatial, and spectral dynamics of brain activity during the resolution of spatial navigation tasks. Our methodological approach also provides proof of concept for future research aiming at singling out oscillatory activity from broadband activity in realistic spatial navigation dynamic tasks. In our opinion, such an analytical approach can help gain new insights into the role of oscillatory brain activity for spatial navigation in humans, as it provides access to the fine temporal structure of brain-behaviour correlations with non-invasive methods.

## ACKNOWLEDGMENTS

This research was supported by grants from the Bial Foundation (Bial 229/20) to M.T.C., and from the *Ministerio de Ciencia e Innovación* (PID2022-137277NB-I00 AEI/FEDER and PRESP05622 PDC2022-133859-I00), and from AGAUR *Generalitat de Catalunya* (2021 SGR 00911) to S.S.F. A.M.M. was partially funded by Maria de Maeztu Units of Excellence Programme CEX2021-001195-M (MICIU/AEI /10.13039/501100011033). We thank Prof. Lluis Fuentemilla for fruitful discussions. The authors acknowledge the invaluable contribution of the subjects who participated in this study.

## ACKNOWLEDGMENTS

This research was supported by grants from Bial Foundation (Bial 229/20) to M.T.C., and from the *Ministerio de Ciencia e Innovación* (PID2022-137277NB-I00 AEI/FEDER and PRESP05622 PDC2022-133859-I00), and from AGAUR *Generalitat de Catalunya* (2021 SGR 00911) to S.S.F. A.M.M. was partially funded by Maria de Maeztu Units of Excellence Programme CEX2021-001195-M (MICIU/AEI /10.13039/501100011033). We thank Prof. Lluis Fuentemilla for fruitful discussions. The authors acknowledge the invaluable contribution of the subjects who participated in this study.

## DATA AVAILABILITY

Pre-processed data and code required to replicate the analysis presented in this manuscript have been uploaded to a private project in Open Science Foundation. Upon acceptance of the manuscript, data, and code will be made publicly available.

## COMPETING INTERESTS

The authors declare no competing interests.

## AUTHORS CONTRIBUTION

Conceptualization: S.S.F. and M.T.C. Data curation: M.T.C. Formal analysis: M.T.C., A.M.M. and M.S.P. Funding acquisition: S.S.F. and M.T.C. Investigation: A.M.M. and M.S.P. Methodology: S.S.F., M.T.C., A.M.M. and M.S.P. Project administration: S.S.F. and M.T.C. Software: M.T.C. Supervision: S.S.F. and M.T.C. Writing-original draft: M.T.C. and S.S.F. Writing-review and editing: M.T.C., S.S.F., M.S.P and A.M.M.

## Footnotes

1 The task that subjects had to perform can be defined as a memory task, as the subjects could solve it by remembering a sequence of turning directions (LRRLR…) whereas other spatial information (like travelled distance, for instance) was not necessary to successfully solve the task. However, in previous publications (Caplan et al., 2001; Kahana et al., 1999) this task has been referred to as a spatial navigation task, so to remain consistent with the literature we will refer to the task as a spatial navigation task.

2 For instance, a theta oscillation at 5 Hz lasting 4 cycles, 800 ms, that started at time 300 ms would not be identified by the method, as only one cycle of the oscillatory activity would fall inside the epoch.

## Notes

### Competing Interest Statement

The authors have declared no competing interest.

### Summary of Updates

This is a revised version of the manuscript and supplementary information.

