## Supplementary Information for "Single-Trial Characterisation of Frontal Theta and Parietal Alpha Oscillatory Episodes during Spatial Navigation in Humans"

*c. Institució Catalana de Recerca i Estudis Avançats (ICREA), Barcelona 08010, Spain*

### 1. CONTROL BEHAVIOURAL ANALYSIS: MAZE PRESENTATION ORDER

To control for a potential effect of maze presentation order on performance, we ran a repeated measures ANOVA with the factors Run (1 to 5) and Order (1 to 5), where Order indicated the order in which the maze was presented during Part II of the experiment, (Delayed Recall mazes). We observed a main effect of Run ($F(4,92)=115.34$, $p<.001$) but no significant effect of Order ($F$ (4,92)=0.58, $p$ =0.7), and no significant interaction between Run and Order ($F$ (4,92)=1.070, $p$ =0.438). Therefore, we conclude that the order of presentation of the mazes did not alter performance.

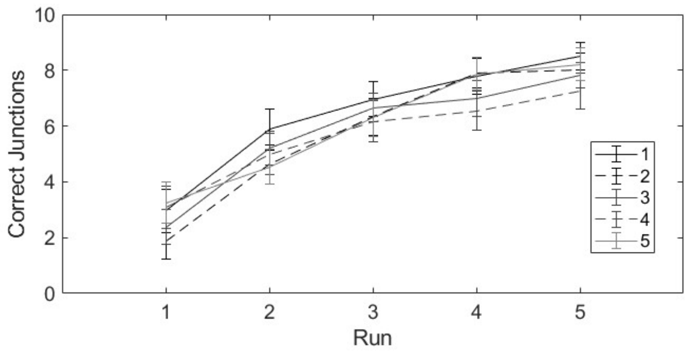


*Supplementary Figure S1: Number of consecutive correct junctions on Delayed Recall mazes as a function of Run. Different lines correspond to the order of presentation of the maze.*

### 2. POWER SPECTRUM AT THE ELECTRODES OF INTEREST

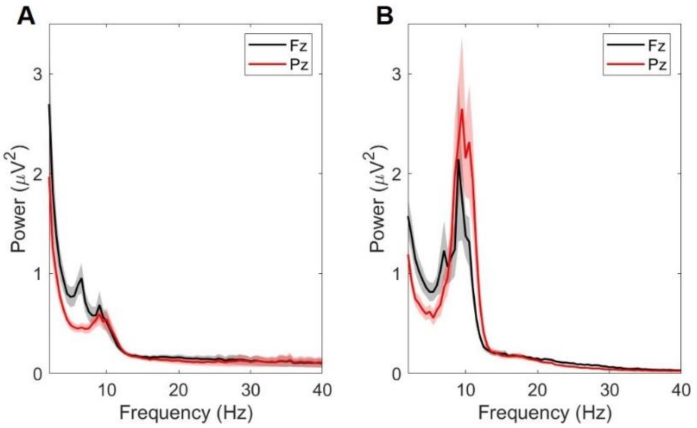


*Supplementary Figure S2: The power spectrum was calculated for the [-6500 -500] ms epochs using Short-Time Fourier Transform, a padding to 8 seconds, Hanning taper at frequencies from 0.5 to 40 Hz in steps of 0.5 Hz. A: Power spectrum during SN task. B: Power spectrum at rest.*

3. INDIVIDUAL DISTRIBUTION OF OSCILLATORY EPISODES

Abundance of oscillatory episodes varied between subjects for both theta and alpha bands (Figure 4). For 19 out of 24 subjects, theta oscillatory episodes were localised in frontocentral areas (Supplementary Figure S3). On the other hand, alpha oscillatory abundance was higher in posterior areas for 17 out of 24 subjects (Supplementary Figure S4).

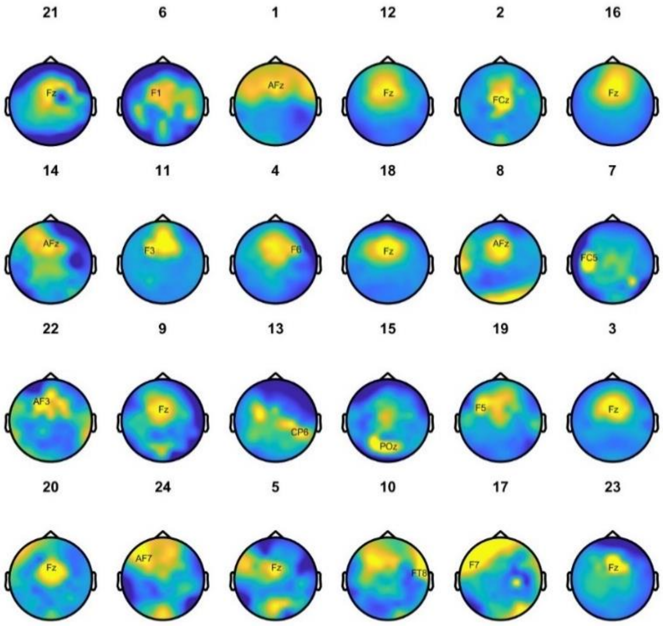


*Supplementary Figure S3: Scalp distribution of the abundance of oscillatory theta episodes for each of the subjects participating in the experiment. Subjects are sorted from maximum to minimum theta abundance. To improve visualization, for each of the subjects the Abundance was z-scored. Dark blue corresponds to z-scored abundance=-2 and bright yellow to z-scored abundance=2.*

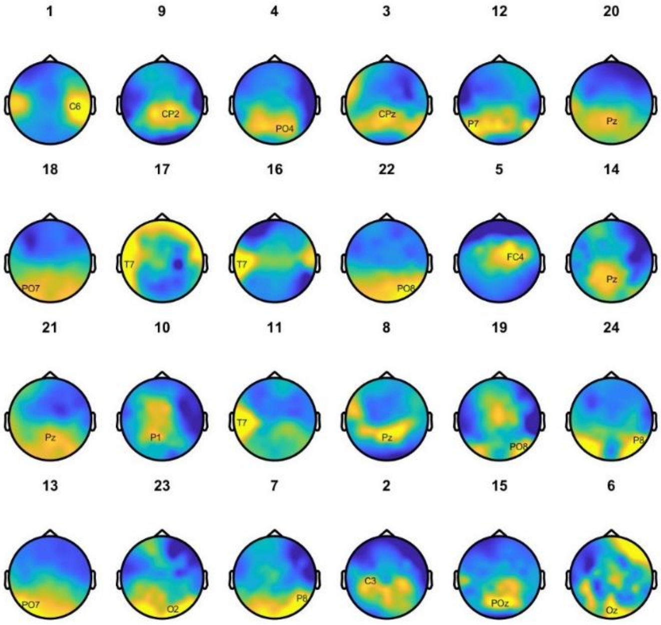


*Supplementary Figure S4: Scalp distribution of the abundance of oscillatory alpha episodes for each of the subjects participating in the experiment. Subjects are sorted from maximum to minimum alpha abundance. To improve visualization, for each of the subjects the Abundance was z-scored. Dark blue corresponds to z-scored abundance=-2 and bright yellow to z-scored abundance=2.*

### 4. LINEAR MIXED MODELS FOR ABUNDANCE

#### 4.1. *Selection of the family*

For each of the frequency bands, the total number of datapoints was 12,720. The dependent variable, Abundance, is a continuous variable in the range [0 1] with a high number of zero values for both theta and alpha band (Supplementary Figure S5). Zero values are informative, as they correspond to instances in which no oscillatory episode was detected so it would be incorrect not to include them in the data. However, it is clear that the data is not normally distributed, and this may result in problems with the fitting.

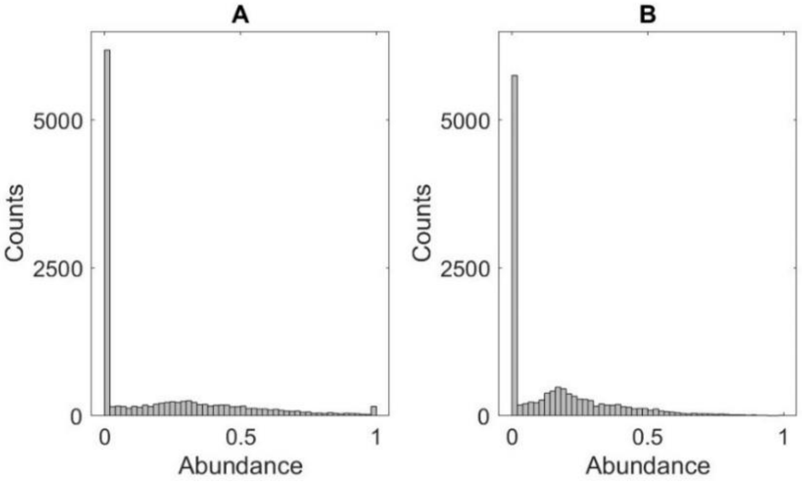


*Supplementary Figure S5: Distribution of Abundance for theta (A) and alpha (B) frequency bands.*

We used the *glmmTMB* (Brooks et al., 2017), R package to fit the full model to our data (for both the theta and alpha frequency band) and we inspected the residuals with *the DHARMa* (Hartig, Florlan, 2018) R package to select the family that best fit our data. We tested gaussian, tweedie (M.C. Tweedie, 1984) and ordered beta (Kubinec, 2023) distributions. For the rest of the tests performed: dispersion, deviation from quantiles, zero inflation, outliers (Supplementary Table 1), ordered beta family provided the best results, with all the tests remaining non-significant except for the outliers in the case of the theta band, and quantiles for the alpha band. Therefore, ordered beta family was selected to fit the model.

#### 4.2. *Model Selection*

As explained in the Methods section, we used the *MuMIn* R package (Bartoń, K., 2009) to rank models according to their second order Akaike Information Criterion (AIC). Additionally, we verified that, for the full model, including the random factor resulted in a reduction of the AIC.

For the theta band, the model that best fit the data was:

$$Abundance \sim Stage + Run + Junction + Time + Correct + (1|Participant)$$

However, this model did not explain the data significantly better than the second-best model (p-value=0.096), with less degrees of freedom. Therefore, we selected the second, simpler, model to explain data in the theta band:

$$Abundance \sim Stage + Run + Junction + Time + (1|Participant)$$

The residuals for the reduced model, similarly to the full model, did not present significant problems (Supplementary Table 2).

For the alpha band, the model that best fit the data across all the models was:

$$Abundance \sim Stage + Junction + Time + Correct + \mathrm{Solved} + \ (1|Participant)$$

However, similarly to the theta band fitting, this model did not significantly improve the fitting of the second-best model (p-value=0.068), that had less degrees of freedom. Therefore, we also selected the simpler, reduced model for the alpha band abundance:

$$Abundance \sim Stage + Junction + Time + Correct + (1|Participant)$$

Like the full model for the alpha band, the deviation from expected quantiles was significant for the reduced model (Supplementary Table 4), although all the other tests performed on the residuals were non-significant.

4.3 *Models for Immediate recall and Delayed recall trials*.

It would be fair to argue that although the task on Immediate and Delayed recall trials is the same, these are, in reality, two very different situations that may rely on different cognitive mechanisms. On Immediate recall trials subjects only required access to short-term memory, with little interference from the other learnt mazes. On the other hand, during Delayed recall trials, subjects had to recall each of the five learnt mazes they were navigating to access information about the correct path. The drop in overall performance for Delayed compared to Immediate recall trials (see *Results* and Figure 2) indicated that the task was significantly easier for Immediate recall trials than Delayed recall trials (two-tailed paired t-test $t(23)=6.84$, $p<.001$, Cohen's d=1.4).

To consider these potential differences, we built separate models for each of the experimental stages (Immediate and Delayed recall) and frequency bands (theta and alpha) following the same procedures as described in the *Statistical analysis* section.

For the theta Band, Immediate recall trials, the model that best described the data was:

$$Abundance \sim Time + Correct + (1|Participant)$$

However, only Correct response was significantly modulating episode abundance (Supplementary Table 3). Post-hoc tests indicated that Abundance of theta episodes was higher on trials in which subjects provided an Incorrect response (Supplementary Table 9). However, it is important to mention that the number of incorrect responses during Immediate Recall mazes was very low (132) compared to correct responses (1597).

For theta band, Delayed recall trials, the model that best described the data was (Supplementary Table 4):

$$Abundance \sim Run + Junction + Time + (1|Participant)$$

Post-hoc tests (Supplementary Table 10) indicated an increase in the abundance of theta episodes on the middle parts of the maze (Junctions 5 and 6) compared to the start of the maze, as well as a decrease in theta episode abundance for the late time window of the trial, -1 second, compared to the early time window of the trial, -5 seconds.

Therefore, it would appear that the model including all trials captures modulations in theta episodes abundance during Delayed recall trials, whereas the differences in theta oscillatory episode abundance between correct and incorrect decisions disappear when all trials are included in the model. A possible interpretation, that must be taken with caution, would be that during the first part of the task (Immediate recall trials), participants had an extremely good performance. On the first run, subjects averaged more than seven Correct decisions in a row, and therefore we can expect that the reliability of the label Correct is high (as the probability of making seven correct choices in a row by chance is 0.8%). On the other hand, for Delayed recall trials, on the first run, subjects took in average of three Correct decisions in a row, we can expect that some of these correct decisions were made by chance (the probability of making three correct choices in a row by chance is 12.5%).

In the alpha band, Immediate recall trials, the model that best described the data did not provide a significant improvement compared to the second-best model ($p=.06$), which had less degrees of freedom. Therefore, we chose the second-best, simpler, model as the one that best fit the data (Supplementary Table 5):

$$Abundance \sim Time + (1|Participant)$$

Post-hoc tests (Supplementary Table 11) indicated a significant decrease in alpha episode abundance in the late time window (-1000 ms) compared to early and middle time windows (-5000 and -3000 ms respectively) of the trial.

Finally, on alpha band, Delayed recall trials, the model that best described the data did not provide a significant improvement compared to the second-best model (*p*=.09), which had less degrees of freedom. Therefore, we chose the second-best, simpler, model as the one that best fit the data (Supplementary Table 6):

$$Abundance \sim Junction + Time + Correct + (1|Participant)$$

Post-hoc tests (Supplementary Table 12) indicated a significant increase in abundance of alpha episodes during the middle parts of the maze (Junctions 3 and 4) as compared to the start (Junction 1) and end (Junction 9) of the maze. In addition, Abundance of alpha episodes was significantly larger for Correct as compared to Incorrect trials.

In summary, the pattern of alpha episodes activity was similar for Immediate recall and Delayed recall trials (Figure 5). Therefore the model considering all the trials provided similar information to the segregated models.

4.4 *Interactions*

All the models described in this paper did not consider interactions between experimental factors. This may result in a loss of relevant information, and therefore, models including interactions were also calculated. Models including interactions were calculated for Immediate and Delayed recall trials separately, because the maximum number of runs on Immediate recall trials was three whereas on Delayed recall trials this was five and this resulted in rank deficient terms in the interaction. In addition, we only included interactions for those terms that appeared in the best model (for each frequency band). Namely, for the theta band we calculated the interactions including Run, Junction, and Time, and for the alpha band we calculated the interactions including Junction, Time, and Correct response. We used the same procedure described in the *Statistical analysis* section (main text) to select the best model. For both frequency bands, the model that best described the data did not include any interactions; hence we retained the models described in the main text.

4.5. *Tables*

4.5.1. Residuals

Supplementary Table S1 includes the results of the tests applied to residuals for the full model for the theta and alpha band for each of the families tested. For each test (dispersion, quantiles, zero inflation, and outliers) the p-value is shown, whereas in the last row, the number of outliers is shown. Ordered beta distribution provides the smaller deviations from expected values for both the theta and alpha band.

| | **Theta** | | | **Alpha** | | |
|---|---|---|---|---|---|---|
| | Gaussian | Tweedie | Ordbeta | Gaussian | Tweedie | Ordbeta |
| **Dispersion** | .8 | .1 | .9 | .9 | .2 | .8 |
| **Quantiles** | <.001 | <.001 | .2 | <.001 | <.001 | .005 |
| **Zero Inflation** | <.001 | .9 | .9 | <.001 | .97 | .97 |
| **Outliers** | <.001 | <.001 | <.001 | <.001 | <.001 | .2 |
| **Number of outliers** | 203 | 61 | 173 | 255 | 52 | 115 |

*Supplementary Table S1: Test of the residuals for models considering all trials (Both Immediate and Delayed recall trials) using different families for modelling Abundance data. Number of subjects: 24, number of observations 12720.*

In Supplementary Table S2, the tests for the residuals of the selected model are shown for each frequency band, all trials (theta and alpha) and Delayed recall only trials (theta DR and alpha DR). As in previous tables all the rows contain p-values but the last row, that contains the number of outliers detected. For all the models, tests for dispersion and zero inflation are non-significant, whereas we found significant deviations from quantiles for model alpha with all trials and alpha Immediate recall only trials and also a significant presence of outliers in all models for theta and the model for alpha Delayed recall only trials, although the maximum percentage of outliers detected was 0.2%.

| | Theta | Alpha | Theta IR | Alpha IR | Theta DR | Alpha DR |
|---|---|---|---|---|---|---|
| **Dispersion** | .96 | .8 | .8 | .9 | .96 | .8 |
| **Quantiles** | .4 | .01 | .91 | .04 | .31 | .09 |
| **Zero Inflation** | .9 | .9~~4~~ | .96 | .9 | .9 | .9 |
| **Outliers** | .001 | .1 | .02 | .2 | .06 | .01 |
| **Number of outliers** | 26 | 18 | 11 | 8 | 13 | 15 |

*Supplementary Table S2: Residual tests for selected models.*

4.5.2 Statistics for the models

Supplementary Tables S3 to S6~~8~~ contain the results of the ANOVA (type III, Wald) for the theta and alpha models for Delayed and Immediate recall trials.

| | $X^2$ | d.f. | p-value |
|---|---|---|---|
| **Participant** | 21.33 | 1 | <.001 |
| **Time** | 5.26 | 2 | .07 |
| **Correct** | 7.82 | 1 | .005 |

*Supplementary Table S3: ANOVA (type III, Wald) for the model describing Abundance in the theta band, Immediate recall only trials.*

| | $X^2$ | d.f. | p-value |
|---|---|---|---|
| **Participant** | 46.81 | 1 | <.001 |
| **Run** | 11.82 | 4 | .02 |
| **Junction** | 34.81 | 9 | <.001 |
| **Time** | 12.39 | 2 | .002 |

*Supplementary Table S4: ANOVA (type III, Wald) for the model describing Abundance in the theta band, Delayed recall only trials.*

| | $X^2$ | d.f. | p-value |
|---|---|---|---|
| **Participant** | ~~84.103~~ | 1 | <.001 |
| **Time** | 29.44 | 2 | <.001 |

*Supplementary Table S5: ANOVA (type III, Wald) for the model describing Abundance in the alpha, Immediate recall only trials.*

| | $X^2$ | d.f. | p-value |
|---|---|---|---|

| | | | |
|---|---|---|---|
| **Participant** | 102.26 | 1 | <.001 |
| **Junction** | 42.66 | 9 | <.001 |
| **Time** | 6.15~~4~~ | 2 | .046 |
| **Correct** | 12.76 | 1 | <.001 |

*Supplementary Table S6: ANOVA (type III, Wald) for the model describing Abundance in the alpha band, Delayed recall only trials.*

4.5.3. Post-hoc tests

For post-hoc tests, we used R package *multcomp* (Hothorn, Torsten et al., 2008), using Tukey contrast and Holm multiple comparison correction. Supplementary Tables S7~~9~~ to S12 contain only the significant contrasts after multiple comparison correction.

| | **Estimate** | **Std. Error** | **z-value** | **p-value** |
|---|---|---|---|---|
| **Delayed recall – Immediate recall** | 0.12 | 0.02 | 4.87 | <.001 |
| **Junction 5 – Junction 1** | 0.21 | 0.04 | 5.32~~4~~ | <.001 |
| **Junction 6 – Junction 1** | 0.16 | 0.04 | 3.98 | .003 |
| **Junction 7 – Junction 1** | 0.15 | 0.04 | 3.47 | .02 |
| **Junction 8 – Junction 1** | 0.15 | 0.05 | 3.36 | .03 |
| **Junction 5 – Junction 2** | 0.13 | 0.04 | 3.25 | .046 |
| **Junction 10 – Junction 5** | -0.23 | 0.06 | -4.15 | .001 |
| **Junction 10 – Junction 6** | -0.19 | 0.06 | -3.29 | .04 |
| **Time -3000 – Time -5000** | -0.06 | 0.02 | -2.56 | .02 |
| **Time -1000 – Time -5000** | -0.09 | 0.02 | -3.96 | <.001 |

*Supplementary Table S7: Theta band, all trials.*

| | **Estimate** | **Std. Error** | **z-value** | **p-value** |
|---|---|---|---|---|
| **Delayed recall – Immediate recall** | 0.05 | 0.02 | 3 | ~~0~~.003 |
| **Junction 3 – Junction 1** | 0.13 | 0.03 | 3.90 | ~~0~~.004 |
| **Junction 4 – Junction 1** | 0.13 | 0.03 | 4.03 | ~~0~~.002 |
| **Time -1000 – Time-5000** | -0.09 | 0.02 | -4.00 | ~~0~~.0001 |
| **Time -1000 – Time -3000** | -0.11 | 0.02 | -5.10 | <.001 |
| **Correct – Incorrect** | 0.12 | 0.03 | 4.29 | <.001 |

*Supplementary Table S8: Alpha band, all trials.*

| | **Estimate** | **Std. Error** | **z-value** | **p-value** |
|---|---|---|---|---|
| **Correct – Incorrect** | -0.16 | 0.06 | -2.80 | .005 |

*Supplementary Table S9: Theta band, Immediate recall trials.*

| | Estimate | Std. Error | z-value | p-value |
|---|---|---|---|---|
| **Junction 5 – Junction 1** | 0.25 | 0.05 | 5.09 | <.001 |
| **Junction 6 – Junction 1** | 0.19 | 0.05 | 3.56 | .016 |
| **Junction 5 – Junction 3** | 0.18 | 0.05 | 3.47 | .02 |
| **Time -1000 – Time -5000** | -0.11 | 0.03 | -3.49 | .001 |

*Supplementary Table S10: Theta band, Delayed recall trials.*

| | Estimate | Std. Error | z-value | p-value |
|---|---|---|---|---|
| **Time -1000 – Time -5000** | -0.13 | 0.03 | -3.95~~1~~ | <.001 |
| **Time -1000 – Time -3000** | -0.18 | 0.03 | -5.23~~0~~ | <.001 |

*Supplementary Table S11: Alpha band, Immediate recall trials.*

| | Estimate | Std. Error | z-value | p-value |
|---|---|---|---|---|
| **Junction 3 – Junction 1** | 0.14 | 0.04 | 3.67 | .010 |
| **Junction 4 – Junction 1** | 0.15 | 0.04 | 3.59 | .014 |
| **Junction 9 – Junction 2** | -0.21 | 0.07 | -3.26 | .04 |
| **Junction 9 – Junction 3** | -0.28 | 0.07 | -4.26 | <.001 |
| **Junction 9 – Junction 4** | -0.29 | 0.07 | -4.25 | <.001 |
| **Correct – Incorrect** | 0.12 | 0.04 | 3.57 | <.001 |

*Supplementary Table S12: Alpha band, Delayed recall trials.*

#### 4.6 *Model selection tables*

In Supplementary Figure S6, the models that account for 99.9% of the total weight of the selection table are displayed. Each column indicates one of the factors, rows correspond to the different models. White cells correspond to factors that are not included in the model of the corresponding row. If a factor is indeed relevant for the data modelled, it should appear consistently in the models that account for most of the weight.

For theta band (Supplementary Figure S6A), Stage, Junction, and Time appear in the models with higher weight, a pattern that is repeated on theta band Delayed recall only trials (Supplementary Figure S6E). However, for theta band Immediate recall (Supplementary Figure S6C) only trials, only Correct response appears consistently among the models with higher weight.

For alpha band (Supplementary Figure S6B), Stage, Junction, Time, and Correct response are the most relevant factors, whereas only Time appears consistently among the models with higher weight for alpha band Immediate recall only trials (Supplementary Figure S6D). Finally, for alpha band, Delayed recall trials (Supplementary Figure S6F), both Junction and Correct response are the most relevant factors.

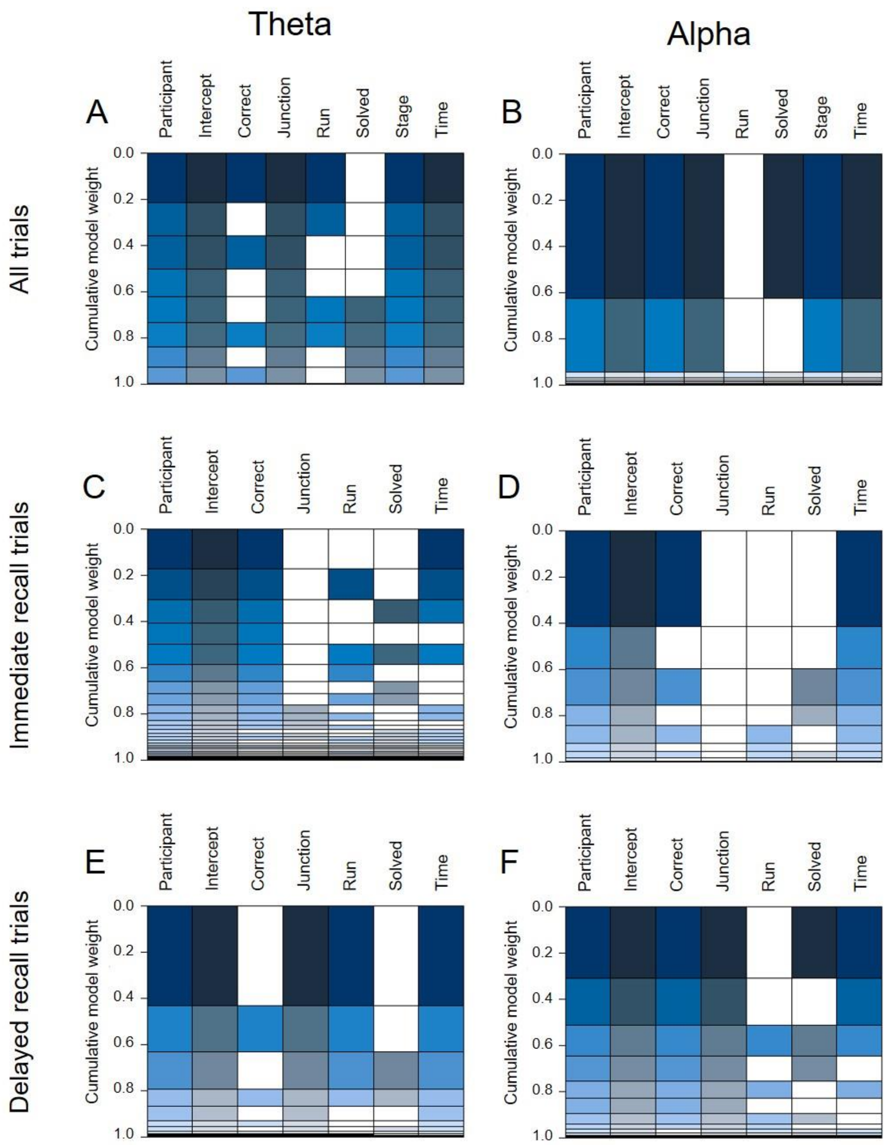


*Supplementary Figure S6: Tables of weights for each of the models. Top row: Models for all trials A: Theta band B: Alpha band. Middle row: Immediate Recall only trials: C: Theta band. D: Alpha band. Bottom row: Delayed recall only trials. E: Theta band. D: Alpha band.*

### 5. LINEAR MIXED MODELS FOR POWER

In addition to models for frontal theta (electrode Fz) and parietal alpha (electrode Pz) abundance, models for power and oscillatory episode power were also calculated for frontal theta and parietal alpha. Power was obtained using the time-resolved spectral estimations generated for the episode detection (see Methods). For the oscillatory episodes power model, power was extracted from the estimation of power obtained with the fBOSC package (Seymour et al., 2022), for time-points were no episode was detected, oscillatory power was set to zero. Both for power and power of oscillatory episodes, equivalent to abundance, time-resolved estimations were averaged for three time windows, 2000 ms duration centered at times -5000, -3000 and -1000 ms relative to the navigational decision. Power and oscillatory episodes power values were averaged across each of the three time-windows of interest.

We evaluated how alpha and theta power varied as a function of task properties. Power at the electrodes displaying maximal activity for each of the frequency bands (Fz for theta and Pz for alpha) was obtained using the spectral estimations generated for the episode detection. These time-resolved power estimations were divided into three time windows, 2000 ms duration centered at times -5000, -3000 and -1000 ms relative to the navigational decision (same time windows as in the main analysis).

For the statistical analysis, the same procedure applied to abundance was followed (see Methods: Statistical Analysis). The distributions that provided the best results for residuals of the power and oscillatory episodes power were lognormal and tweedie (M.C. Tweedie, 1984).

5.1 Models for frontal theta power and episode power.

The results for both power and oscillatory episode power for frontal theta (3 to 8 Hz) were in line with the results for frontal theta abundance. For theta power, the model that best described the data was (see Supplementary Table S13):

$$Power \sim Stage + Run + Junction + Time + (1 \dashv |Participant)$$

Post-hoc tests indicated that Theta power was significantly higher during Delayed recall mazes as compared to Immediate recall mazes, on Run 2 as compared to Run 1, on Junctions 5 to 7 as compared to Junction 1, on Junctions 5 and 6 as compared to Junction 10, during time window -5000 ms as compared to -3000 and -1000 and during time window -3000 as compared to -1000.

| | $X^2$ | d.f. | p-value |
|---|---|---|---|
| **Participant** | 123440.53 | 1 | <.001 |
| **Stage** | 29.88 | 1 | <.001 |
| **Run** | 13.41 | 4 | .01 |
| **Junction** | 127.91 | 9 | <.001 |
| **Time** | 253.85 | 2 | <.001 |

*Supplementary Table S13: ANOVA (type III, Wald) for the model describing power in the theta band.*

For oscillatory episode power, the model that best described the data was (see Supplementary Table S14):

$$Power \sim Stage + Run + Junction + Time + Correct + (1 \dashv |Participant)$$

Although the factor Correct did not contribute significantly to the model (Supplementary Table S14). Post hoc tests indicated increased power of theta episodes for Delayed recall trials as compared to Immediate recall trials, on Run 4 as compared to Runs 1 to 3, on Junctions 3 to 6 and Junction 8 as compared to Junction 1, on Junction 5 as compared to Junctions 9 and 10, during Time window -5000 ms as compared to Time windows -3000 and -1000 ms and during Time window -3000 as compared to -1000 ms.

| | $X^2$ | d.f. | p-value |
|---|---|---|---|
| **Participant** | 1264.39 | 1 | <.001 |
| **Stage** | 9.19 | 1 | .002 |
| **Run** | 16.55 | 4 | .002 |
| **Junction** | 50.49 | 9 | <.001 |
| **Time** | 26.06 | 2 | <.001 |
| **Correct** | 3.73 | 1 | .053 |

*Supplementary Table S14: ANOVA (type III, Wald) for the model describing episode power in the theta band.*

In summary, for theta episodes, the results show a robust and consistent pattern that associates more abundant and stronger theta oscillatory episodic activity with access to spatial information during navigation.

5.2 Models for parietal alpha power and episode power.

The model that best described parietal alpha power differed from the one obtained for the episodes. The model that best represented the data was:

$$Power \sim Run + Correct + Solved + (1 \,|Participant)$$

But this model was not significantly better than the second best, more parsimonious model:

$$Power \sim Run + Solved + (1 \,|Participant)$$

Therefore, we selected the simpler model to represent the data (Supplementary Table S15). Post-hoc tests, indicated an increase in alpha power on Run 4 as compared to Runs 1 and 2 and on Solved mazes as compared to Unsolved mazes.

| | $X^2$ | **d.f.** | **p-value** |
|---|---|---|---|
| **Participant** | 3706.72 | 1 | <.001 |
| **Run** | 17.44 | 4 | .002 |
| **Correct** | 2.65 | 1 | .1 |
| **Solved** | 9.37 | 1 | .002 |

*Supplementary table S15: ANOVA (type III, Wald) for the model describing power in the alpha band.*

For parietal alpha oscillatory episodes power, the model that best described the data included all the factors (Supplementary Table S16):

$$Power \sim Stage + Run + Junction + Time + Correct + Solved + (1 \,|Participant)$$

Post hoc tests indicated that the power of alpha episodes was significantly higher on Runs 3 and 5 as compared to Run 1, on Junction 3 as compared to Junctions 1 and 2 and to Junctions 5 to 8, during Time windows -5000 and -3000 ms as compared to -1000 ms, on trials preceding a Correct response, and on mazes that were Solved vs. Unsolved.

In summary, switches to internal attention can be related to increased parietal alpha episodic activity that reflects an increased abundance and power of episodic activity on correct as compared to incorrect trials, and during specific moments of the task. However, other factors (as Solved compared to Unsolved mazes) present increased episodic oscillatory power but not abundance. We believe that more research, out of the scope of current study, should be performed before providing an interpretation.

| | $X^2$ | **d.f.** | **p-value** |
|---|---|---|---|
| **Participant** | 272.72 | 1 | <.001 |
| **Stage** | 3.56 | 1 | .06 |
| **Run** | 15.33 | 4 | .004 |
| **Junction** | 46.05 | 9 | <.001 |
| **Time** | 60.97 | 2 | <.001 |
| **Correct** | 6.23 | 1 | .01 |
| **Solved** | 9.48 | 1 | .002 |

*Table S16: ANOVA (type III, Wald) for the model describing episode power in the alpha band.*

6. STEERING WHEEL SPECIFICATIONS

The steering wheel used was a Logitech G29, compatible with Windows® systems 7, 8, 8.1 10, and 11 and Play Station 3,4, and 5. The system consisted of a steering wheel (900º rotation lock to lock, two motor force feedback, Hall effect steering sensor, and overheat safeguard) and a pedal panel (three pedals, nonlinear brake pedal, patented carpet grip system, textured heel grip, and self-calibrating system)."

7. CYBERSICKNESS

7.1 *Online screening of subjects susceptible to suffering cybersickness*

All potential participants had to fill out the Cybersickness Susceptibility Questionnaire (CSQ) (Freiwald et al., 2020), as part of an online screening process, prior to scheduling their experimental sessions. The subjects that were cleared to participate in the experiment had a mean score of 0.7±0.48 [0.1 – 1.7] on the Motion Sickness portion of the questionnaire, which corresponded to "rarely" experiencing motion sickness in response to the questionnaire items. In contrast, the nine individuals that were advised against participating due to a heightened likelihood of experiencing cybersickness had a mean score of 1.7±0.40 [1.3 – 2.3] on the motion sickness subscale, or a rating of occasionally experiencing motion sickness. Particular interest was given to the items referring to cars (items 1, 2) and buses (items 7, 8) given their similarity to the experimental environment. The excluded participants also showed higher overall ratings on these four items of interest (Supplementary Figure S7 and Supplementary Table S17).

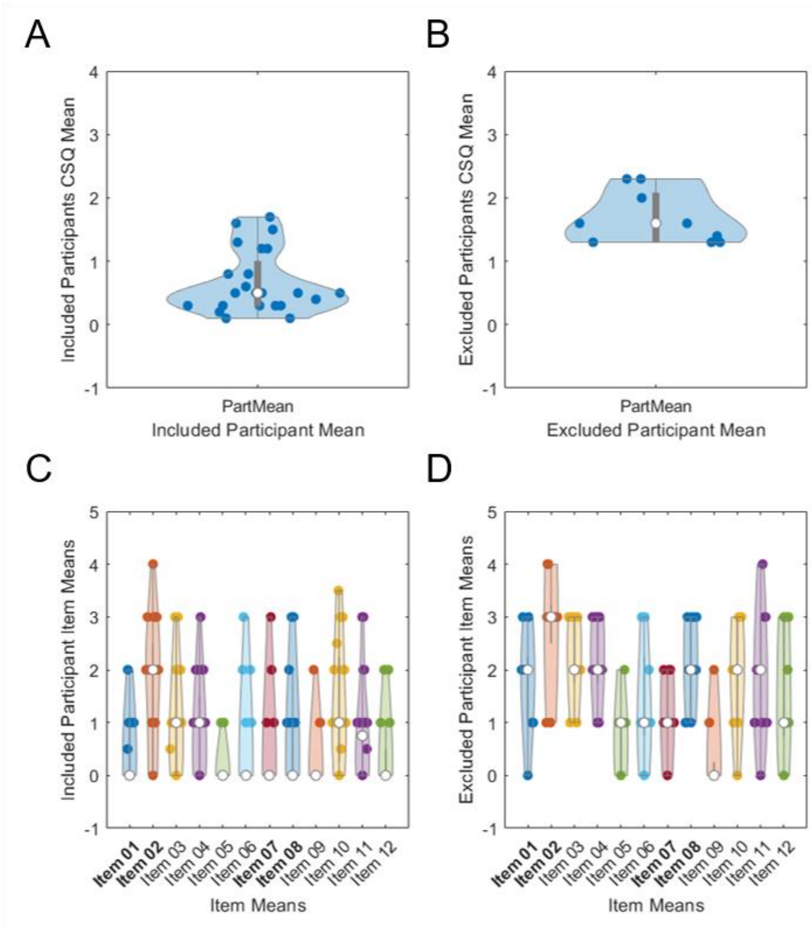


*Supplementary Figure S7: **Top**: Scores on the Cybersickness Susceptibility Questionnaire (CSQ) (Freiwald et al., 2020) averaged across items for included (**A**) and excluded (**B**) subjects. **Bottom**: Scores for individual items of the CSQ for included (**C**) and excluded (**D**) subjects. Coloured dots correspond to individual values and white dots to the average across subjects. In panels **C** and **D**, the items in bold are the ones that were more relevant to the online screening process.*

| | Item 1 (When I drive a car, I feel sick as a passenger) | Item 2 (Reading as a passenger makes me sick) | Item 7 (I feel sick when I ride the bus or am a co-driver in a car) | Item 8 (I feel sick when I ride the bus sitting backwards) |
|---|---|---|---|---|
| Included Participants | 0.4±0.65 | 1.6±1.24 | 0.3±0.75 | 0.5±0.93 |
| Excluded Participants | 1.8±0.97 | 2.8±1.09 | 1.2±0.67 | 2.1±0.93 |

*Supplementary Table S17: Average scores for the four items of interest of the CSQ* (Freiwald et al., 2020) *for subjects included and excluded in the experiment.*

7.2 *Screening of subjects experiencing adverse symptoms before entering VR environment*

Upon arrival, subjects were screened for the presence of potentially adverse symptoms (with particular interest being placed on headache, blurred vision, and eye strain) using the Simulator Sickness Questionnaire (Kennedy et al., 1993). None of the subjects reported the presence of severe symptoms (mean Total Score: 2.0±1.57 [0 – 5.0] calculated using the Bouchard et al. (2007, 2021) unweighted scoring approach to avoid inflating total scores) on any of the SSQ items. The maximum Total Score could reach a value of 60, meaning that participants were well below this maximum prior to immersion in the virtual reality environment. In terms of the items of interest, mean raw scores were 0 (headache), 0.1±0.41 (blurred vision), and 0.2±0.38 (eye strain), well below the maximum of 3. No subjects were excluded based on the results of this questionnaire, given that none of them reported feeling severely affected by any of the items pre-immersion (Supplementary Figure S8).

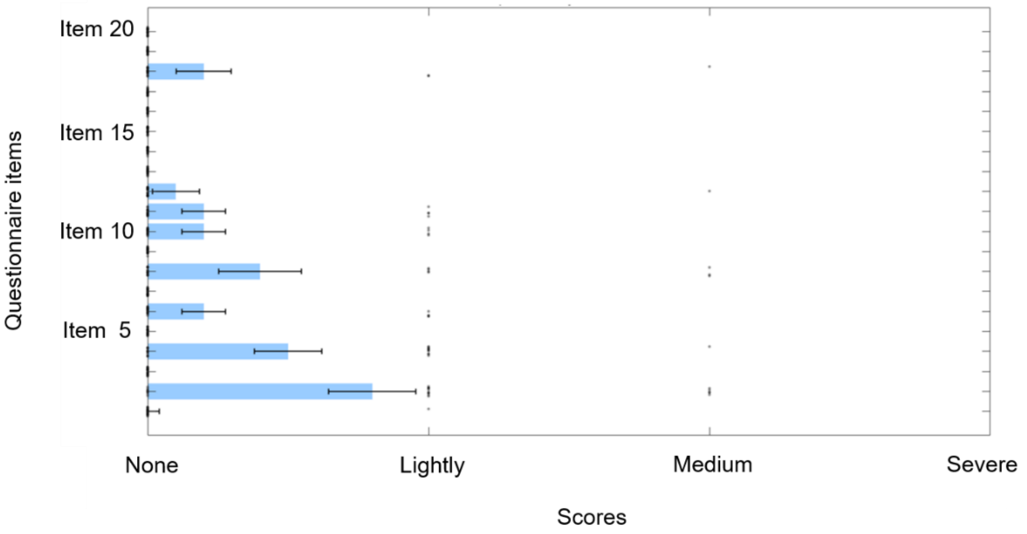


*Supplementary Figure S8: Scores for the items of the Simulator Sickness Questionnaire (Kennedy et al., 1993) provided by the subjects before entering the virtual reality environment. Blue bars indicate average across subjects, error bars correspond to standard deviation and black dots correspond to individual responses.*
